# Diet-induced obesity dysregulates chromatin oxygen sensing regulating efferocytosis in macrophages

**DOI:** 10.1101/2023.05.12.540252

**Authors:** Kentaro Takahashi, Jinghua Liu, Jasmine R. Jackson, Muthusamy Thiruppathi, Elizaveta V. Benevolenskaya, Timothy J. Koh, Norifumi Urao

**Affiliations:** Department of Pharmacology, State University of New York Upstate Medical University, Syracuse, NY 13210, USA; Department of Kinesiology and Nutrition, University of Illinois at Chicago, Chicago, IL 60612, USA; Department of Biochemistry and Molecular Genetics, University of Illinois at Chicago, Chicago, IL 60612, USA

**Keywords:** obesity, high-fat diet, macrophages, metabolism, histone modification, hematopoietic stem cells, hypoxia, oxygen, oxidative stress

## Abstract

Macrophages are plastic cell populations that normally adapt to their environment. Cellular adaptation to hypoxia occurs through transcription factors including hypoxia-inducible factors, and hypoxia-inducible transcriptions are further regulated by chromatin response through histone modification including histone methylation. However, the role of histone methylation in the hypoxia response of macrophages is not well understood. As obesity is associated with dysregulated macrophage functions, we investigated whether hypoxia response is cell-intrinsically dysregulated in macrophages in obesity.

In mouse bone marrow-derived macrophages (BMDMs), immunoblotting revealed that 1% hypoxia rapidly increases the global levels of histone 3 methylations. We found that hypoxia-induction of histone 3-lysine 4 tri-methylation (H3K4me3) is specifically inhibited in BMDMs from mice fed a high-fat diet (HFD-BMDMs) compared to BMDMs from mice fed a normal diet (ND-BMDMs). Multi-omics approach with ChIP-seq and RNA-seq identified that glycolysis-related pathways and genes including *Aldoa* are upregulated after prolonged hypoxia along with upregulated H3K4me3 in ND-BMDMs. In contrast, no pathway is associated with hypoxia-upregulated H3K4me3 peaks in HFD-BMDMs and hypoxia-induced *Aldoa* expression is decreased in HFD-BMDMs, suggesting both the extent and the genome location of H3K4me3 response to hypoxia is dysregulated in obesity. Consistently, lactate accumulation and induction of histone lactylation under hypoxia are reduced in HFD-BMDMs. Furthermore, HFD-BMDMs exhibited decreased dying cell clearance under hypoxia due to the reduced capacity of anaerobic glycolysis. Competitive bone marrow transplantation of hematopoietic stem cells (HSCs) shows that HFD-induced long-term memory reflects the impaired dying cell clearance in differentiated BMDMs, which is rescued by inhibiting oxidative stress in HSCs.

In summary, chromatin response to hypoxia associated with H3K4me3 enrichment governs transcriptions for anaerobic glycolysis and dying cell clearance under hypoxia. Obesity dysregulates the extent and the genome location of H3K4me3 enrichment, glycolysis, and dying cell clearance of BMDMs under hypoxia, which is initiated in HSPCs via oxidative stress.

## Introduction

Macrophages are ubiquitous in mammals and humans, involving tissue homeostasis and pathologies. Both systemic and local inflammation increase the contributions of blood monocyte-derived macrophages to the total inflammatory or immune cell populations, determining whether the inflamed tissue will be recovered through the resolution of inflammation or will be unresolved and sustained inflammation ^1^. Macrophages are highly plastic for their phenotypes, as the most simply described as inflammatory (M1-like) and homeostatic (M2-like) phenotypes, which have a corresponding metabolic signature—glycolysis and oxidative phosphorylation, respectively ^2^. Recent studies showed that macrophages equip an internal clock for their phenotypic transition from inflammatory to homeostatic phenotype based on the intracellular lactate build-up resulting from hypoxia or pro-inflammatory stimuli^3^, highlighting cell-intrinsic mechanisms that ensure macrophage plasticity in the normal state.

Obesity is a major health problem causing its associated conditions such as type 2 diabetes mellitus, hypertension, and coronary heart diseases^4^. Obesity dysregulates macrophage functions such as phagocytosis^5^ and their phenotypic plasticity^6^, which have been observed as an increased number of inflammatory macrophages resulting in poor prognosis of tissue healing in ischemic diseases or injured sites. Systemic metabolic conditions can undergo reprogramming, which in turn changes bone marrow-derived monocyte/macrophage functions and epigenetic signatures in their precursor cells ^7^. It remains to be understood how obesity dysregulates intracellular machinery promoting macrophage plasticity.

Hypoxia governs important physiological and pathological processes ^8^. In homeostatic conditions, cells in most organs reside in an environment with lower oxygen levels than ambient levels, which have circadian oscillation^9, 10^. In pathological conditions, especially with ongoing ischemia or inflammation, cells are often exposed to hypoxia due to disruption of the vasculature or higher demand for oxygen in inflammatory cells. Hypoxia-inducible factors (HIFs) are essential oxygen-sensing elements that regulate their target genes to exercise cellular adaptation to lower oxygen such as angiogenesis, glycolysis, or cell death^11, 12^. As a transcriptional activity of HIF-dependent and –independent target genes is influenced by the chromatin state—such as open, poised, and closed chromatin, oxygen-sensitive chromatin modifications provide additional layers of transcription regulation under hypoxia^13–16^. It has been shown that hypoxia induces histone hypermethylation ^17^, as Jumonji C (JmjC) containing histone lysine demethylases (KDMs) contain prolyl hydroxylase in their structure, and thus act as molecular hypoxia sensors that directly modify histone methylation, resulting in either activation or suppression of gene expression ^17^.

Dying cell clearance by macrophages, which is typically studied as efferocytosis, a phagocytic process clearing apoptotic cells, promote pro-healing programs in and nearby macrophages^18^, in which glycolysis and lactate production play crucial roles in maintaining efferocytotic capability^19^. Moreover, macrophages primed by prolonged hypoxia activate the transcription of glycolytic genes, which is associated with promoted efferocytosis capacity^20^. However, whether the metabolic changes that increase dying cell clearance involve chromatin remodeling by histone modifications remains to be understood.

Here, we identify oxygen-sensing chromatin response by H3K4me3 in macrophages that potentially regulate the broad range of metabolic pathways using the multi-omics approach, which in turn promotes lactate accumulation, reduced oxidative stress, and dying cell clearance under hypoxia. In macrophages from diet-induced obese mice, this oxygen-sensing H3K4me3 response is largely dysregulated, leading to impaired dying cell clearance capacity under hypoxia. We found that the transcription levels of the glycolytic enzyme, *Aldoa*, under prolonged hypoxia, is associated with the rapidly upregulated H3K4me3 at its promoter under hypoxia. Moreover, supplementing glucose or lactate is sufficient to increase the defective dying cell clearance capacity in obese macrophages. As a top of the cascade events of hypoxia response, the oxygen sensing H3K4me3 response finetunes the metabolic changes such as glucose utilization and glycolysis, which specifically prime macrophages for promoted dying cell clearance. Given that dying cell clearance is critical for the resolution of inflammation, targeting the H3K4me3 response may provide a novel therapeutic strategy for non-resolving inflammation in obesity.

## Methods

### Animals

All animals were housed in a temperature-controlled facility with a 12-h light/dark cycle in compliance with Upstate Medical University Institutional Animal Care and Use Committee protocols. C57BL/6 male mice at the age of 8 weeks were purchased from The Jackson Laboratory (Bar Harbor, ME) and fed a high-fat diet (HFD, 60% kcal fat, Research Diets Inc. New Brunswick, NJ, USA) or normal diet (ND, 10% kcal fat, Research Diet Inc.) for 16-20 weeks.

### Cells

Bone marrow-derived macrophages (BMDMs) were isolated and cultured as previously described. Briefly, bone marrow (BM) cells were collected from femur and tibia bones with centrifugation at 10,000g for 15 sec. After resuspended with culture media, cells were seeded in plates and cultured in RPMI medium with 10% heat-inactivated FBS (ThermoFisher, Waltham MA, USA) and 20 ng/ml recombinant mouse macrophage colony-stimulating factor (M-CSF, PeproTech, Cranbury, NJ, USA, 315-02). After 3 days of incubation at 37℃ with 5% CO_2_, the media was replaced with fresh medium containing M-CSF. Cells were allowed to differentiate into macrophages for a total of 7 days and then were harvested by removing the supernatant. BMDMs were exposed to 1% hypoxia in the hypoxia chamber (BioSpherix, C-174 and ProOx C21, Parish, NY, USA) for 1 hour or 24 hours before harvesting or performing assays. Bone marrow-derived dendritic cells were generated by culturing bone marrow progenitors in RPMI medium supplemented with 10% FBS and 20ng/ml granulocyte-macrophage colony-stimulating factor (GM-SCF, PeproTech) for 10 days.

### Immunofluorescence imaging

For immunofluorescence imaging, 1.2 × 10^5^ BMDMs were cultured on a chamber slide (ThermoFisher). Cells were fixed with 4% paraformaldehyde (Electron Microscopy Sciences, Hatfield, PA, USA) for 10 minutes, permeabilized with 0.2% Triton X-100 (Millipore Sigma, Burlington, MA, USA) for 10 minutes, blocked with 1% donkey serum (Jackson ImmunoReseach Inc., West Grove, PA, USA) for 30 minutes, and incubated with rabbit anti-H3K4me3 antibody (Abcam, ab8580, Cambridge, UK) overnight. Cells were then washed in PBS and incubated with 1:1000 anti-rabbit AlexaFluor 647-conjugated secondary antibody (ThermoFisher) for 30 minutes. After washing with PBS, the cells were stained with Hoechst 33258 (Millipore Sigma). Images were obtained with an automated cell imaging system (ImageXpress Pico, Molecular Devices, San Jose, CA, USA) and high-content imaging analysis was performed to count cells and quantify signal intensity with CellReporterXpress (Molecular Devices).

### ChIP-seq library preparation and sequencing

ChIP-seq experiments were performed with the rabbit anti-H3K4me3 antibody (Abcam) in BMDMs derived from HFD mice or ND mice after incubation in 1% hypoxia treatment or normoxia for 1 hour. Chromatin samples were prepared using the Zymo-Spin ChIP kit (Zymo Research, Irvine, CA, USA) according to the manufacturer’s instructions. Briefly, BMDMs were crosslinked with 0.4% paraformaldehyde for 10 minutes at room temperature (RT) and immediately quenched with 0.125 M glycine (Millipore Sigma) for 5 minutes at RT. For the hypoxia treatment group, samples were processed in the large hypoxia chamber (BioSpherix, C-Shuttle and ProOx P360) to maintain the hypoxic environment during procedures. After washing with PBS containing 1 mM phenylmethylsulfonyl fluoride (PMSF, Millipore Sigma) and cOmplete EDTA-free protease inhibitor cocktail (Millipore Sigma), BMDMs were resuspended in 500 µl chilled nuclei prep buffer and incubated on ice for 5 minutes. 2 × 10^6^ BMDMs were sonicated using a Covaris M220 sonicator (Covaris, Woburn, MA, USA) at a high power setting (peak power: 150, duty factor: 20, cycles: 200) for 330 seconds to yield a modal fragment size of 150 bp. After incubation with 1 µg H3K4me3 antibody at 4℃ overnight, ChIP-DNA was prepared using ZymoMag Protein A beads (Zymo Research). Input samples were also prepared using 10% of chromatin samples. ChIP and input libraries were prepared using NEBNext Ultra II DNA Library Prep Kit for Illumina (New England Biolabs, Ipswitch, MA, USA). Libraries were sequenced on an Illumina NextSeq500 (Illumina, San Diego, CA, USA) with High Output NextSeq kit (Illumina) using 2x 75bp paired-end reads.

### ChIP-seq data processing

The fastq files were processed using PartekFlow (Partek Inc., St. Louis, MO, USA). Briefly, the raw reads were aligned to the mouse genome annotation mm10 with Bowtie2. Peaks were called by MACS2 peak caller in narrow peak mode with the IP sample bam files set as treatment and corresponding input bam files set as control^21^. Called peaks were quantified to regions and the normalization was performed with transcripts per million (TPM), and the normalized peaks were then subjected to statistical analysis using Partek gene-specific analysis (GSA) algorithm. Differential H3K4me3 peaks were identified with thresholds (p<0.05, least square [LS] mean>20).

Promoter peaks were defined as ±3kb from the nearest transcription start site (TSS), intergenic peaks were defined as >3kb upstream of the nearest TSS and >3kb of the nearest transcription end site (TES), and gene body peaks were defined as all peaks not identified as promoter or inter genic peaks.

To visualize the coverage tracks, bam files were merged with Samtools merge, and merged bam files were normalized with counts per million (CPM) followed by conversion to bedgraph files with bamCoverage. Merged bedgraph files were visualized with the IGV genome browser. Pathway enrichment analysis was performed using the Molecular Signatures Database (MSigDB) online tool provided by Gene Set Enrichment Analysis (GSEA) with hallmark genes and <0.05 p-value cut-off^22, 23^. ChIP-sequencing data is deposited under NCBI GEO accession GSE231991.

### RNA isolation

Cells were lysed with TRI Reagent (Zymo Research), and total RNA was extracted with Direct-zol RNA Miniprep Kit (Zymo Research) according to the manufacturer’s instructions. RNA quantity was evaluated with QubitFlex and Qubit broad-range RNA assay kit (ThermoFisher). The quality of RNA samples was assessed with the Agilent 2100 Bioanalyzer (Agilent Technologies), and all samples showed RNA integrity number > 9.

### RNA-seq library preparation and sequencing

RNA sequencing libraries were prepared using the Illumina Stranded Total RNA Prep with Ribo-Zero Plus (Illumina). Once prepared indexed cDNA libraries were pooled in equimolar amounts and sequenced with single-end 75bp reads on an Illumina NextSeq500.

### RNA-seq data processing

The fastq files obtained from RNA-seq and GSE192969 were processed using PartekFlow (Partek Inc., St. Louis, MO, USA). In brief, the raw reads were aligned using STAR and the aligned reads were quantified to the annotation model through Partek E/M. The normalization was performed with the median ratio through PartekFlow. The normalized counts were subjected to statistical analysis using GSA. Differentially expressed genes (DEGs) were identified with thresholds (p<0.05, fold change<-2.0 or 2.0<fold change)^22, 24^. RNA-sequencing data was deposited under NCBI GEO accession GSE231992.

### Gene ontology analysis

Gene ontology (GO) analysis for differential peaks and differentially expressed genes was performed using Molecular Signatures Database (MSigDB, https://www.gsea-msigdb.org/gsea/index.jsp)^22,24^ or Ingenuity Pathway Analysis (QIAGEN, Germantown MD, US). P value<0.05 was considered statistically significant.

### Quantitative RT-PCR

1,000 ng of RNA was reverse transcribed using iScript (BioRad, Hercules, CA, USA) and qPCR was performed using iTaq Universal SYBR Green Supermix (BioRad) in duplicates. Ct values were obtained on CFX Opus 384 Real-Time PCR System (BioRad), and data was generated with the comparative threshold cycle (Delta CT) method by normalizing to hypoxanthine phosphoribosyltransferase (*Hprt*).

### Immunoblots

BMDMs were lysed with M-PER™ Mammalian Protein Extraction Reagent (ThermoFisher) containing cOmplete™ Mini Protease Inhibitor Cocktail (Millipore Sigma) as per the manufacturer’s instructions. Lysate was centrifuged for 10min at 14,000g at 4℃, and supernatant was collected. For extraction of histones, BMDMs were washed with pre-warmed serum-free RPMI and lysed with ice-cold extraction buffer (Active Motif, Carlsbad, CA, USA) with cOmplete™ mini protease inhibitor cocktail (Millipore Sigma). Cells were homogenized with vigorous pipetting and cell lysate was centrifuged to obtain the crude histone. A neutralization buffer (Active Motif) containing 0.1M DTT (Millipore Sigma) and a Halt protease inhibitor cocktail (ThermoFisher) was added to the crude histone. Protein concentration was quantified using Qubit™ Protein and Protein Broad Range (BR) Assay Kits (ThermoFisher).

25µg of protein lysates were denatured at 95℃ for 5 minutes in 4x Laemmli Sample Buffer (BioRad) containing β-Mercaptoethanol (Millipore Sigma). Protein samples were separated on 4-15% SDS-PAGE gels (BioRad) at 100V and electro-transferred to 0.45µm nitrocellulose membranes (BioRad) at 50V for 85min (for cytoplasmic protein) or to 0.2µm nitrocellulose membranes (BioRad) at 30V for 70min (for histones). The membranes were blocked with EveryBlot Blocking Buffer (BioRad) for 5 min and incubated for 1h at room temperature with primary antibodies. The appropriate HRP-conjugated secondary antibodies were used for the chemiluminescent detection of proteins with Clarity Max Western ECL Substrate (BioRad). Membranes were scanned with a ChemiDoc imaging system (BioRad) and quantified using Image Lab 6.1.0 software (BioRad). The following primary antibodies were used for immunoblotting: Hif1α (36169, Cell Signaling), Arg1 (GTX109242, GeneTex), β-actin (3700, Cell Signaling), H3K4me3 (ab8580, Abcam), H3K9me3 (13969, Cell Signaling), H3K27me3 (9733, Cell Signaling), H3K18la (PTM-1406, PTM BIOLABS), and total H3 (3638, Cell Signaling). The following secondary antibodies were used for immunoblotting: anti-rabbit HRP-linked IgG antibody (5196-2504, BioRad) and anti-mouse HRP-linked IgG antibody (7076, Cell Signaling).

### Induction of apoptosis in neutrophils and Jurkat cells

Murine neutrophils were isolated with density gradient centrifugation-based protocol, as previously described^25^. Briefly, bone marrow cells harvested from mouse femurs and tibias were centrifuged with Histopaque 1119 (Millipore Sigma) and Histopaque 1077 (Millipore Sigma). Neutrophils were collected at the interface of Histopaque 1119 and Histopaque 1077. A mixture of early/late apoptosis and some necrotic death was induced by irradiating Jurkat cells and the isolated neutrophils under a UV lamp (254nm, Stratalinker UV 1800, Stratagene Corporation, La Jolla, CA) for 1 and 15 min, respectively. The cells were incubated in the CO_2_ incubator for 3 hr. This method routinely yielded greater than 80% Annexin V-positive cells including early-and late-stage apoptosis.

### Dying cell clearance assay

BMDMs were plated in a 96-well plate at a density of 0.4 × 10^5^ cells per well and exposed to hypoxia or normoxia for 24 h prior to the assay. Apoptotic neutrophils or Jurkat cells were stained with CellTracker™ Red CMTPX Dye (ThermoFisher) by incubating for 20 min, and BMDMs were stained with carboxyfluorescein succinimidyl ester (CFSE, ThermoFisher) by incubating for 20 min. Labeled dying cells were added to BMDMs at a 5:1 ratio and co-cultured for 1 h followed by vigorous washing 3 times with 1x PBS to remove dying cells. Cells were fixed with 1% paraformaldehyde (Electron Microscopy Sciences) and imaged with an automated cell imaging system (ImageXpress Pico, Molecular Devices) and analysis was performed to count CellTrackerRed^+^/CSFE^+^ BMDMs with CellReporterXpress (Molecular Devices).

For live cell imaging, dying cells were stained with 1.3µM pHrodo Green STP Ester (ThermoFisher) by incubating for 45min at 37℃ after apoptosis induction. Labeled dying cells were added to BMDMs at a 5:1 ratio and cells were imaged with Image Xpress Pico in the environmental control unit which introduces normoxia or 1% hypoxia at 37℃ and 5% CO_2_ environment. Images were taken every 10min for up to 6 hours with ImageXpress Pico. Signal intensity was measured with CellReporterXpress.

For continual dying cell clearance assay, the first round of dying cells was labeled with CellVue Claret Far Red (Millipore Sigma) and co-cultured with macrophages at a 1:1 ratio for 45min.

Then, apoptotic cells were washed away using cold assay media. Macrophages were subsequently rested in the CO_2_ incubator for 2h prior to co-culture with a second round of dying cells that were labeled with pHrodo Green. Continual efferocytosis was assessed by live cell imaging as CellVueClaret^+^ (1st dying cell uptake) and CellVueClaret^+^ pHrodoGreen+ (2nd dying cell uptake) BMDMs. Images were taken every 10min for up to 2 hours with ImageXpress Pico. Signal intensity was measured with CellReporterXpress.

### Lactate measurement

Intracellular lactate concentration was quantified using Lactate-Glo Assay (Promega, Madison, WI, USA) after lysing 50,000 BMDMs with 0.6 N HCl (Millipore Sigma) containing 0.16% Dodecyl trimethylammonium bromide (Millipore Sigma) according to manufacturer’s instructions. Luminescence was measured with a Synergy H1 plate reader (Agilent Technologies).

### Analysis of cellular metabolic activity

BMDMs were seeded into XFe96 Microplates (Agilent Technologies, Santa Clara, CA, USA). After 24-hour culture, the medium was changed to Seahorse XF base medium (Agilent Technologies) supplemented with 5 mM glucose, 2 mM glutamine, and 1 mM sodium pyruvate, and the BMDMs were incubated for 1 hour in the no CO_2_ incubator. The extracellular acidification rate (ECAR) was measured using the Seahorse XF96e analyzer (Agilent Technologies) under basal conditions and after administration of mitochondrial inhibitors (0.5 µM rotenone/antimycin A, Agilent Technologies) and 2-deoxy-d-glucose (50 mM 2-DG, Agilent Technologies) as a glycolysis inhibitor. The glycolytic proton efflux rate (PER) was calculated in the WAVE software (Agilent Technologies).

### Hematopoietic stem and progenitor cell isolation

Cell surface staining and hematopoietic stem and progenitor cell (HSPC) isolation procedures were performed as described previously ^26^. In brief, bone marrow (BM) cells were collected from femur and tibia bones from transgenic mice expressing EGFP (C57BL/6-Tg(CAG-EGFP)131Osb/LeySopJ, Jackson Laboratory) with centrifugation at 10,000g for 15 sec, and purified by Ficoll separation with Histopaque-1119 (Millipore Sigma). Cells were stained with unconjugated rat anti-mouse lineage-specific antibodies (Ter-119, Mac1, Gr-1, B220, CD5, CD3, CD4, CD8, and CD127, Biolegend, San Diego, CA, USA), followed by incubating with Dynabeads anti-rat IgG (Thermo Fisher). Unbound cells including HSPCs were collected after magnetic sorting. Cells were then stained with goat anti–rat PE-Cy5 secondary antibody (Thermo Fisher) for lineage-specific primary antibodies, and c-kit–APCeFluor780 (Thermo Fisher), Sca1-PB (Biolegend), CD150-PE (Biolegend) antibodies. Cell sorting was performed on a FACS AriaIII cell sorter (BD). Data analysis was performed using FlowJo v10.8 software (BD). In the cyclosporin A treated group, HSPCs were isolated in buffers containing 50 µg/ml.

### Oxidative stress level measurement

BMDMs or HSPCs were stained with 5 µM CellROX Green (ThermoFisher) for overall oxidative stress measurement or 10 µM MitoPY1 (Biotechne, Minneapolis, MN, USA) for mitochondrial oxidative stress measurement by incubating at 37℃ for 30min. BMDMs were detached from the plates by incubating in D-PBS at 4℃ for 30min after staining. Stained cells were analyzed with a flow cytometer (Cytek Aurora: Cytek Biosciences, Fremont, CA, USA) and FlowJo v10.8 (BD Biosciences, Franklin Lakes, NJ, USA).

### Competitive bone marrow transplantation

For competitive transplantation assays, 8–12-wk-old C57BL/6 recipient mice were lethally irradiated (9 Gy in one dose) and retroorbitally injected with 5,000 purified SLAM HSCs (EGFP+) together with 100,000 whole BM cells obtained from C57BL/6 mice (EGFP-) as competitors. Peripheral blood was collected by retroorbital bleeding at 8 and 16 weeks after transplantation and stained with specific antibodies for flow cytometry analysis. For serial transplantation, HSPCs were isolated 16 weeks after the first transplantation. Lethally irradiated 8-12-week-old C57BL/6 recipient mice were injected with 3,000 HSCs (EGFP+) together with 100,000 whole BM cells obtained from C57BL/6 mice (EGFP-) as competitors. BMDMs were isolated 16 weeks after 2^nd^ transplantation and a dying cell clearance assay was performed with pH-sensitive dye CypHer5E NHS Ester (1 µM, Cytiva, Marlborough, MA USA) with/without hypoxia treatment.

## Results

### Hypoxia rapidly increases histone methylation in macrophages

Histone methylation can be induced in response to hypoxia in multiple cell types, in acute manners^15^. To test if such hypoxia-induced histone methylations are preserved in macrophages, we used primary murine macrophages differentiated from macrophage precursors in the bone marrow (BMDMs). Fully differentiated BMDMs were exposed to hypoxia (1% O_2_) for 1 hour, and we analyzed global levels of trimethylation of histone H3 lysine 4 (H3K4me3), trimethylation of histone H3 lysine 9 (H3K9me3), and trimethylation of histone H3 lysine 27 (H3K27me3) using western blotting. As shown in the control BMDMs obtained from mice fed a normal-fat diet (ND-BMDM), significant upregulation of H3K4me3 levels after 1-hour hypoxia, together with other histone methylations such as H3K9me9 and H3K27me3 (Fig. 1a,b). The immunofluorescence high-content imaging confirmed that global levels of H3K4me3 were increased in ND-BMDM after 1-hour hypoxia (Fig. 1c,d).

**Figure 1:**
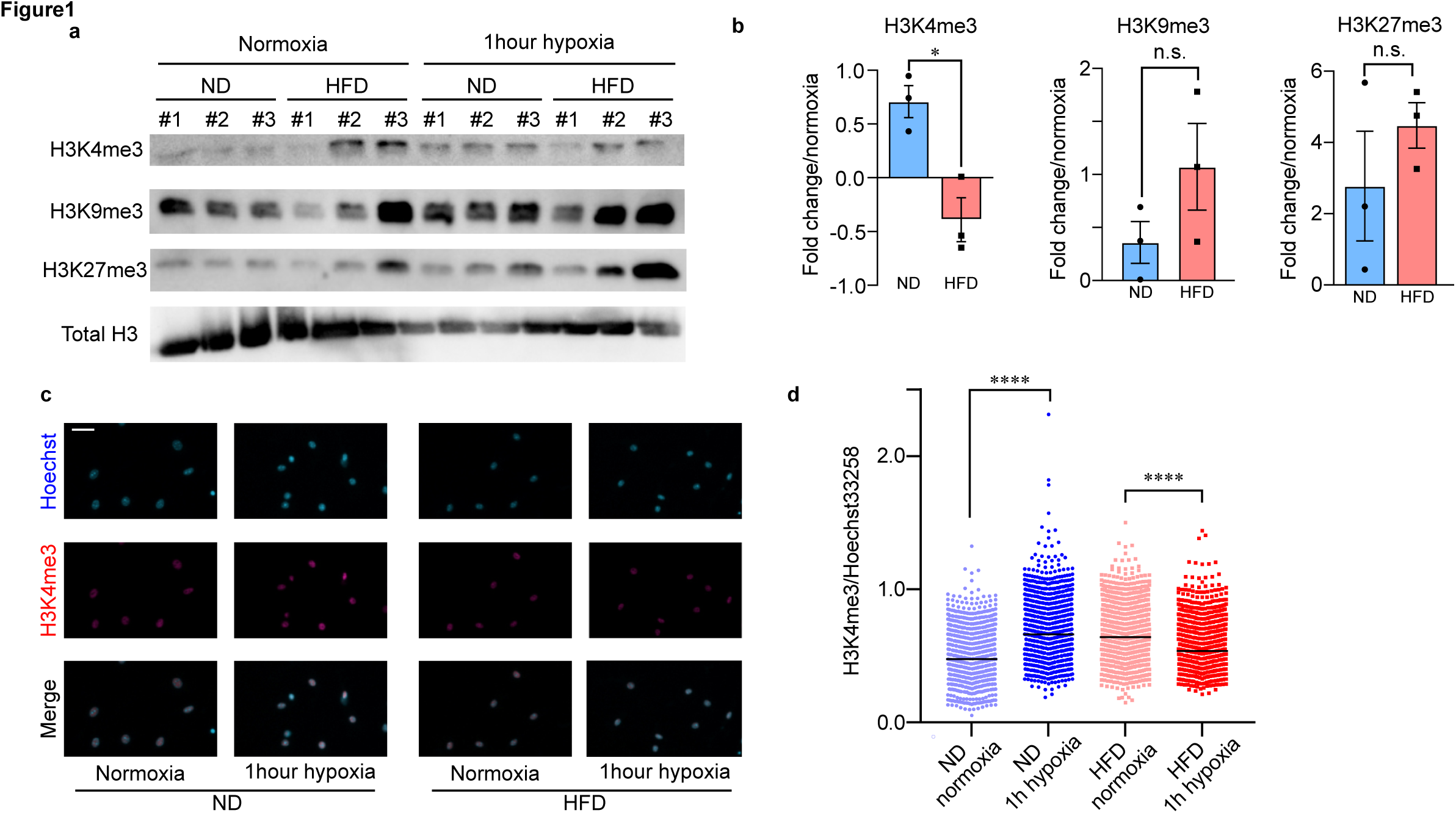
Obesity impairs hypoxia-inducible histone methylation in bone marrow-derived macrophages (BMDMs) a: Immunoblots of acid-extracted histones for histone methylation after hypoxia treatment for 1 hour. ND: BMDMs obtained from normal diet mice, HFD: BMDMs obtained from high-fat diet mice. b: Hypoxia-induced change of histone methylation. The relative enrichment of histone methylation versus total H3 was quantified by densitometry. Three biological replicates from n=3 mice per group; statistical significance was determined by Student’s t-test. c: High content imaging of H3K4me3 in BMDMs after 1hour hypoxia. BMDMs were stained with Hoechst33258 (Blue, top) and H3K4me3 antibody (Red, middle). [Bottom] Merged images with H3K4me3 and Hoechst33258. The scale bar indicates 100 µm. d: Ratio of H3K4me3/Hoechest33258 intensity analyzed by high-content imaging. n=7000-8000 cells obtained from three biological replicates; statistical significance was determined by two-way ANOVA with Turkey’s HSD post hoc test. All values are means ± SEM; * p<0.05, * * * * p<0.0001; n.s. not significant.

### Obesity impairs rapid hypoxia-induced H3K4me3 enrichment in macrophages

We investigated how obesity affects the global levels of histone methylations in macrophages in response to hypoxia. To do this, we tested BMDMs obtained from mice fed a high fat-diet (HFD-BMDMs) and found that hypoxia failed to induce H3K4me3 enrichment in HFD-BMDMs, while hypoxia-induced H3K9me3 and H3K27me3 enrichment were comparable to ND-BMDMs (Fig 1a,b). The high-content imaging recapitulated the impaired H3K4me3 response to hypoxia in HFD-BMDMs (Fig. 1c,d). In the same hypoxic condition, the levels of HIF-1α were similarly increased in both ND-and HFD-BMDMs (Supplementary Fig. 1), suggesting that diet-induced obesity impairs rapid hypoxia-induction of H3K4me3 in macrophages in a HIF-1-independent manner.

### Hypoxia-upregulated H3K4me3 peaks mark metabolic genes enriched under prolonged hypoxia in macrophages

Next, we performed ChIP-seq to identify the genomic location of H3K4me3 peaks modified by a brief (1 h) exposure to hypoxia (1% O_2_). Among the total of 25,247 peaks in ND-BMDMs (Supplementary Fig. 2), we found 698 differential peaks after the brief hypoxia treatment — upregulated (448 peaks, 64.2%) or downregulated (250 peaks, 35.8%) — compared to normoxia (Fig. 2a, Table 1, p-value < 0.05, LSmean ≥ 20). Consistent with a previous report in a different cell type^15^, more than 80% of hypoxia-upregulated peaks were located at promoters of gene locus as shown in *Epb41l4b* locus, while a small part of the differential peaks is located at either intergenic region or gene body (Fig. 2b,e). By assigning a differential peak to a gene whose transcription starting site (TSS) is closest to the peak, we found that multiple metabolic pathways including glycolysis, oxidative phosphorylation, and fatty acid metabolism are identified by Gene Set Enrichment Analysis (GSEA) of hypoxia-upregulated H3K4me3 peaks at promoter regions (398 peaks) (Fig. 2b, and Table 2).

**Figure 2:**
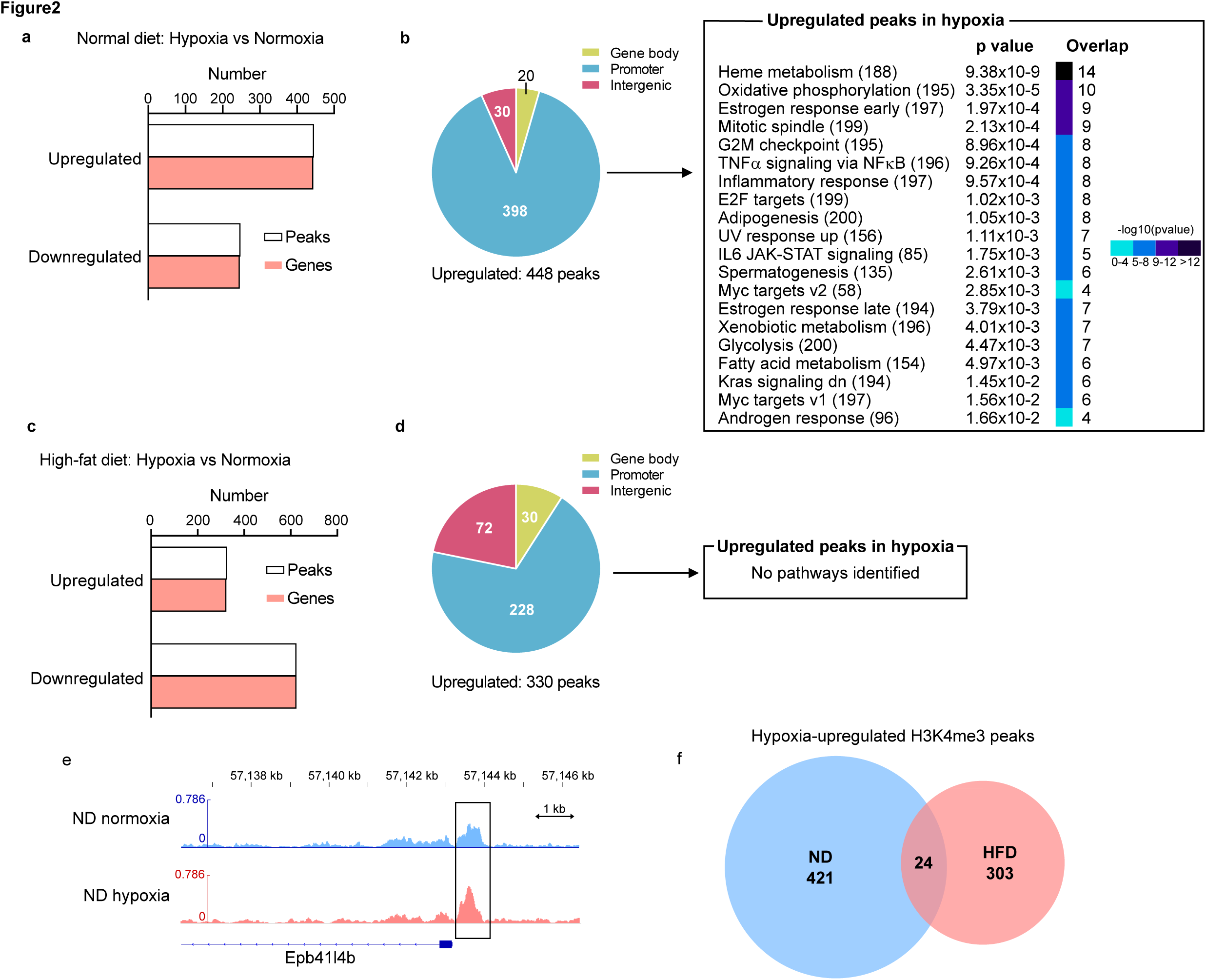
Characterization of hypoxia-inducible H3K4me3 peaks in ND-BMDMs and HFD-BMDMs. a: The number of differential H3K4me3 peaks after the brief hypoxia treatment in ND-BMDMs. b. Distribution of upregulated H3K4me3 peaks on gene locus and associated pathways identified by gene ontology (GO) analysis in ND-BMDMs. [Left] The pie chart illustrates the location of differential H3K4me3 peaks around the annotated genes. [Right] GO analysis revealed significant enrichment of gene set signatures for upregulated H3K4me3 peaks after the brief hypoxia treatment. c. The number of differential H3K4me3 peaks after the brief hypoxia treatment in HFD-BMDMs. d. Distribution of upregulated H3K4me3 peaks on gene locus and associated pathways identified by gene ontology (GO) analysis in HFD-BMDMs. [Left] The pie chart illustrates the location of differential H3K4me3 peaks around the annotated genes. [Right] GO analysis failed to show pathways associated with upregulated H3K4me3 peaks in HFD-BMDMs. e. Coverage tracks of normalized H3K4me3 peak at *Epb41l4b* locus. Hypoxia-induced H3K4me3 peak at the promoter region. f. Overlaps of hypoxia-upregulated H3K4me peaks between ND-BMDMs and HFD-BMDMs.

We took a multi-omics approach to examine the impact of the hypoxia-modified H3K4me3 peaks on transcription in BMDMs. For transcription, we performed RNA-seq analysis in our cultured BMDMs as well as analyzed available datasets of bone marrow-derived immortalized macrophages primed with prolonged hypoxia (GSE192969) ^20^. As hypoxia-induced genes are increased over time from 1 hour to 7 days of hypoxia exposure, the overlapping hypoxia-upregulated H3K4me3 peaks with gene transcripts as well as with associated pathways are increased (Fig. 3a and 3b). Notably, glycolysis and fatty acid metabolism, but not oxidative phosphorylation, were identified as hypoxia-upregulated metabolic pathways marked by H3K4me3 upregulation (Fig. 3c). At the individual gene level, we identified *Aldoa* and *Depdc1a* as hypoxia-increased glycolytic genes marked by hypoxia-upregulated H3K4me3 peaks (Fig. 3d). As we did not find overlaps between hypoxia-upregulated H3K4me3 peaks and hypoxia-increased genes at 1-hour hypoxia (Fig. 3b), the rapidly upregulated H3K4me3 peaks likely induce the poised chromatin state that influence transcriptional activation under prolonged hypoxia.

**Figure 3:**
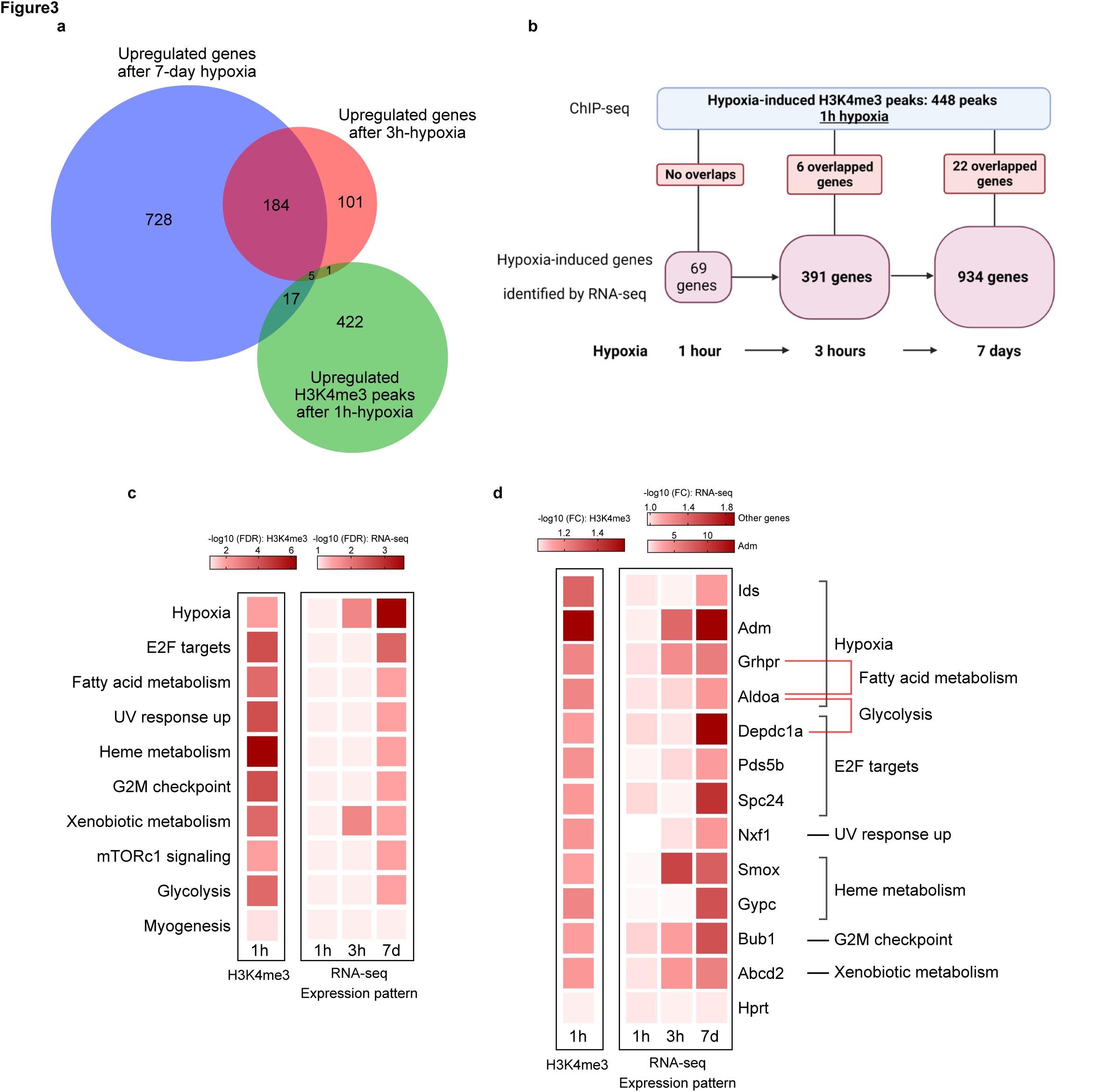
Hypoxia-upregulated H3K4me3 peaks after brief hypoxia overlap genes enriched under prolonged hypoxia in macrophages. a: Venn diagram demonstrates overlapping genes among enriched genes after 3-hour– and 7-day– hypoxia, and hypoxia-upregulated H3K4me3 peaks after 1-hour-hypoxia. b. Time course of overlapping genes between upregulated H3K4me3 peaks and enriched gene expression after hypoxia treatment. c. Gene ontology (GO) terms enriched among genes overlapping between hypoxia-upregulated H3K4me3 peaks after brief hypoxia treatment and enriched genes after a brief and prolonged hypoxia treatment. d. Gene expression pattern of overlapping genes between hypoxia-upregulated H3K4me3 peaks after brief hypoxia treatment and enriched genes after a brief and prolonged hypoxia treatment.

### Obesity dysregulates hypoxia-upregulated H3K4me3 peaks and metabolic gene enrichment under prolonged hypoxia in macrophages

As obesity impairs rapid H3K4me3 upregulation by hypoxia measured as protein levels (Fig. 1a), we then investigated the genomic location of H3K4me3 peaks in HFD-BMDMs by ChIP-seq, which identified a total of 958 differential peaks including 330 upregulated and 628 downregulated peaks (Fig. 2c) — 118 less upregulated and 378 more downregulated by hypoxia compared to ND-BMDMs. Among 330 upregulated H3K4me3 peaks in HFD-BMDMs, 170 less at promoter regions and 42 more at intergenic regions are found than ones in ND-BMDMs, and GSEA demonstrated that hypoxia-upregulated H3K4me3 peaks in HFD-BMDMs are not associated with any metabolic pathways that were shown in ND-BMDMs (Fig. 2d). Only 24 peaks are the overlap of hypoxia-upregulated H3K4me3 peaks between ND– and HFD-BMDMs (Fig. 2f). Thus, obesity-induced dysregulation in hypoxia-upregulated H3K4me3 peaks spans broadly from the number of peaks to the genomic locations.

We next examined the impact of this lack of proper H3K4me3 response to hypoxia in HFD-BMDMs on transcription by focusing on metabolic genes. As we identified *Aldoa* and *Depdc1a* as hypoxia-increased glycolytic genes marked by hypoxia-upregulated H3K4me3 peaks in the multi-omics approach (Fig. 3d), we quantitatively analyzed these genes using qPCR analysis. We confirmed that HFD-BMDMs lack *Aldoa* enrichment in prolonged hypoxia (24 h), which was observed in ND-BMDMs, while there was no difference in *Depdc1a* (Fig. 4e). Among 22 genes that are potentially regulated by hypoxia-upregulated H3K4me3 (Fig. 3d), we found *Adm* is also enriched by prolonged hypoxia in ND-BMDMs, which is dampened in HFD-BMDMs (Fig. 4f). To study acute transcriptional change after hypoxia (1 hour), which might be affected by the lack of proper H3K4me3 response to hypoxia in HFD-BMDMs, we explored RNA-seq data in ND– and HFD-BMDMs. In terms of hypoxia-enriched genes, the gene sets including hypoxia, mTORC1 signaling, and glycolysis were associated both in ND– and HFD-BMDMs (Supplemental Fig. 4c,5c) with minimum differences in individual gene repertoire (Supplemental Fig. 4f). To investigate the more detailed expression pattern in response to hypoxia, we further categorized differential genes into four groups (Supplementary Fig. 6a). In Group 1, 53 genes with the enrichment dampened by HFD; in Group 2, 260 genes with the enrichment enhanced by HFD; in Group 3, 145 genes with the reduction enhanced by HFD; and in Group 4, 18 genes with the reduction dampened by HFD. In both Group 1 and Group 2, which represent the enrichment by hypoxia either dampened or enhanced by HFD, Hypoxia, mTORC1 signaling, Glycolysis, and TNFα signaling via NFκB are listed as the top four gene sets (Supplementary Fig. 6b). We did not see a clear link between differentially regulated H3K4me3 peaks and transcripts, either. These data suggest that acute enrichment of genes by hypoxia is not impaired in HFD-BMDMs and that the lack of proper H3K4me3 response to hypoxia in HFD-BMDMs does not affect acute transcription of hypoxia-responsible genes, supporting the claim that hypoxia-upregulated H3K4me3 induces poised chromatin state for later activation of transcription.

**Figure 4:**
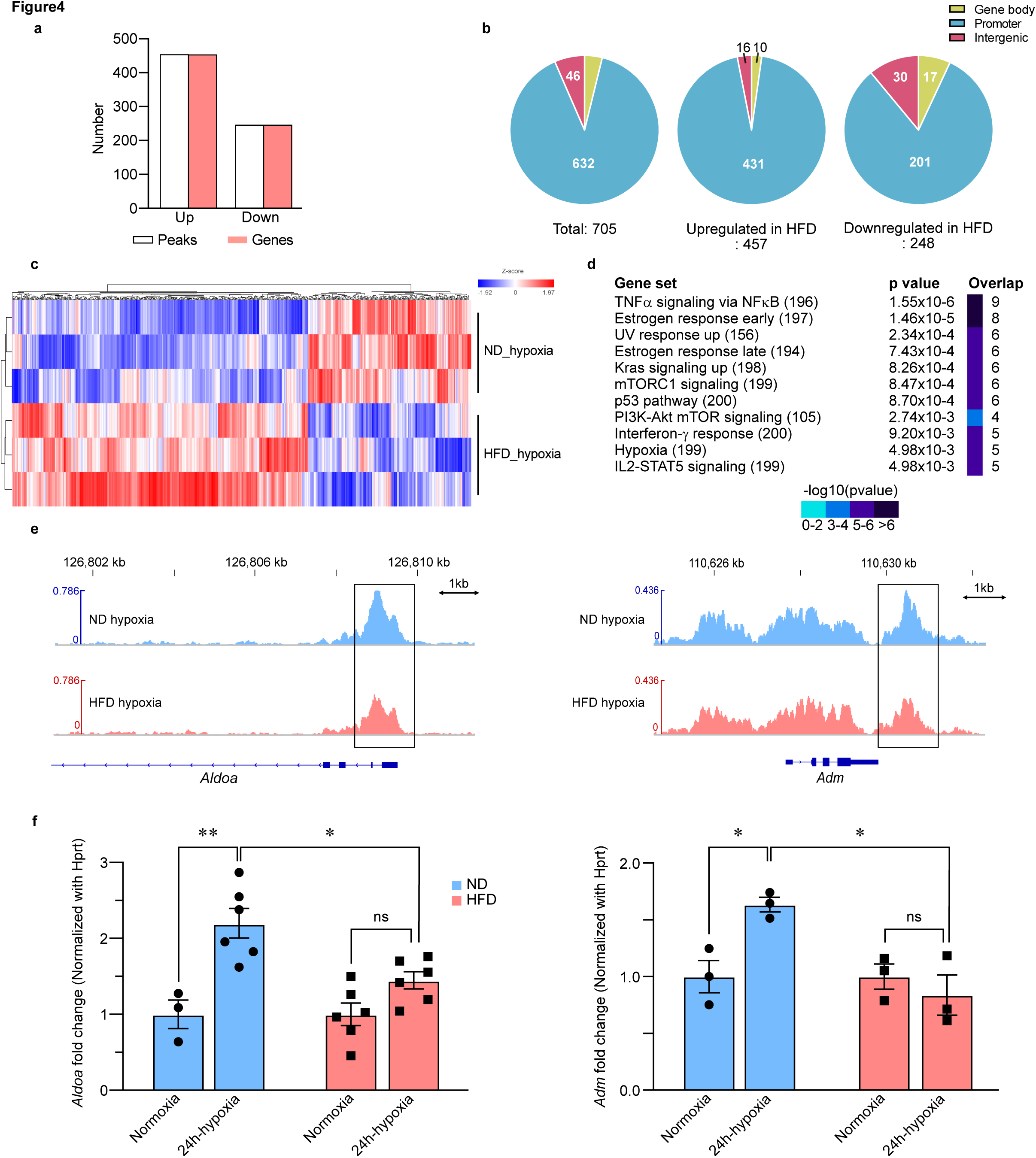
Obesity impairs hypoxia-upregulated H3K4me3 peaks in metabolic genes. a: The number of differential H3K4me3 peaks in HFD-BMDMs over ND-BMDMs after the brief hypoxia treatment. n=3 biological replicates. b. Distribution of upregulated and downregulated H3K4me3 peaks on gene locus. The pie chart illustrates the location of differential H3K4me3 peaks around the annotated genes. c. Heat map of 705 differential H3K4me3 peaks in HFD-BMDMs. The red and blue color indicates increased and decreased H3K4me3 in HFD-BMDMs, respectively. d. GO analysis for 248 HFD-downregulated H3K4me3 peaks in HFD-BMDMs shows the association with signaling pathways including hypoxia. e. Coverage tracks of normalized H3K4me3 peak at *Aldoa* and *Adm* locus in hypoxia condition. H3K3me peaks at promoter regions were downregulated in HFD-BMDMs compared to ND-BMDMs. f. Gene expression of *Aldoa* and *Adm* in ND-BMDMs and HFD-BMDMs after prolonged hypoxia. The rate of hypoxia induction was assessed by normalizing the relative expression level in hypoxia with the one in normoxia. n=3-6 biological replicates; statistical significance was determined by two-way ANOVA with Turkey’s HSD post hoc test. All values are means ± SEM; * p<0.05, * * p<0.01; n.s., not significant.

**Figure 5:**
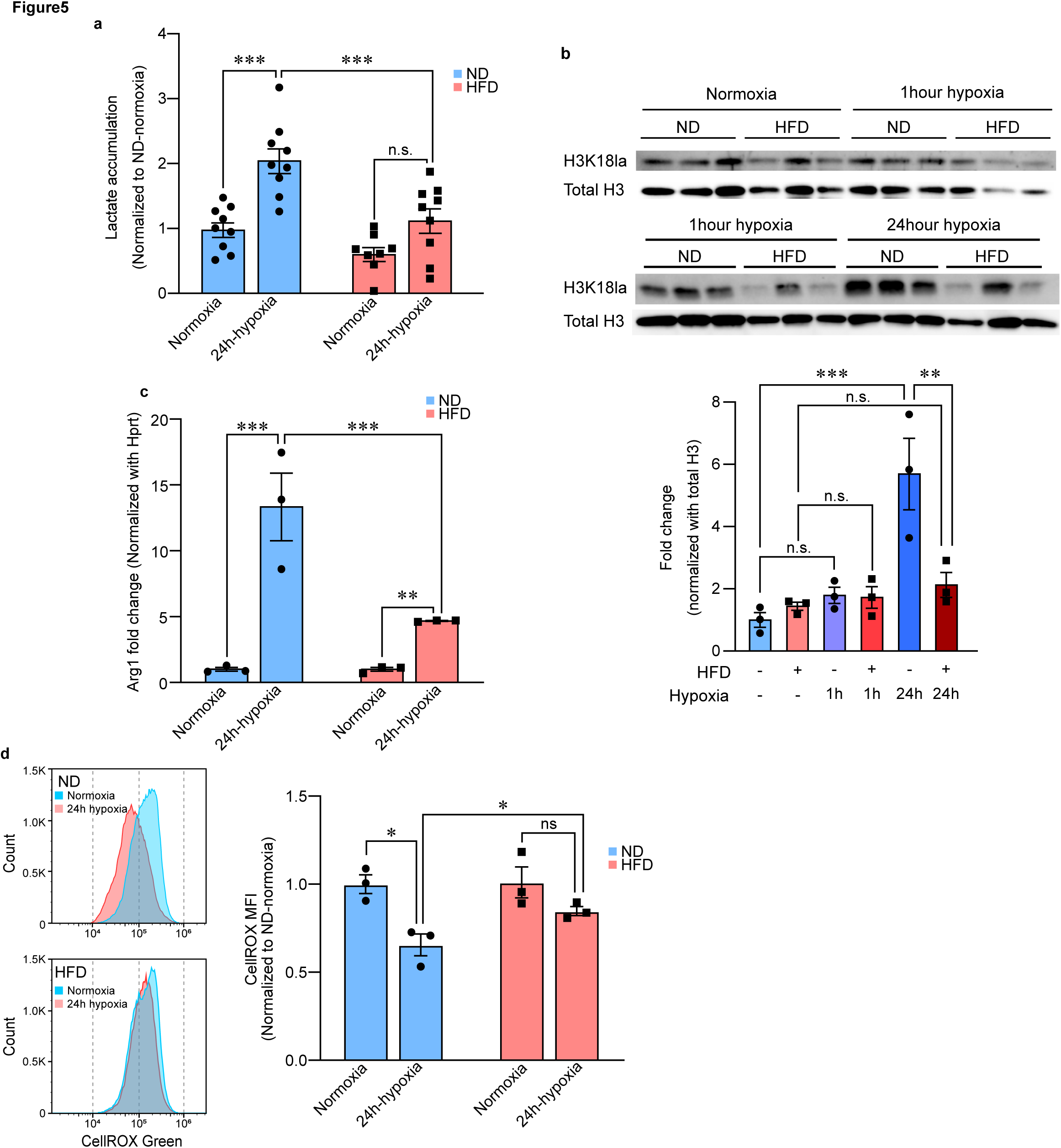
Obesity-dysregulated hypoxia-inducible metabolic change. a. Lactate accumulation in BMDMs after prolonged hypoxia treatment. n=9 biological replicates; statistical significance was determined by two-way ANOVA with Turkey’s HSD post hoc test. b. Histone lactylation enrichment in response to hypoxia. [Top] Immunoblots of acid-extracted histones from BMDMs in response to a brief and prolonged hypoxia. [Bottom] The relative enrichment of H3K18la versus total H3 was quantified by densitometry. n=3 biological replicates; statistical significance was determined by one-way ANOVA with Turkey’s HSD post hoc test. c. Hypoxia-inducible gene expression of *Arg1* in ND-BMDMs and HFD-BMDMs. The rate of hypoxia induction was assessed by normalizing the relative expression level in hypoxia with the one in normoxia. n=3 biological replicates; statistical significance was determined by two-way ANOVA with Turkey’s HSD post hoc test. d. Oxidative stress levels in BMDMs were assessed by measuring the CellROX indicator with flow cytometry. [Left] Representative flow cytometry histograms measuring CellROX Green in BMDMs with or without hypoxia treatment. [Right] Measured CellROX signal intensity was normalized with mean fluorescence intensity (MFI). n=3 biological replicates; statistical significance was determined by two-way ANOVA with Turkey’s HSD post hoc test. All values are means ± SEM; * p<0.05, * * p<0.01, * * * p<0.001; n.s., not significant.

### Obesity dysregulates metabolic changes induced by hypoxia

We next studied metabolic states in BMDMs. A Seahorse assay did not identify alteration of glycolytic capacity in the steady state in the atmospheric oxygen (Supplementary Fig. 7). To measure glycolytic capacity after prolonged hypoxia (for 24 hours) in 1% O_2_ environment in which extracellular acidification rate does not properly reflect glycolysis with the Seahorse assay, we measured lactate accumulation in BMDMs. ND-BMDMs accumulated lactate intracellularly under prolonged hypoxia, whereas HFD-BMDMs did not show lactate accumulation under prolonged hypoxia (Fig. 5a). Moreover, histone lactylation, a recently defined histone modification as a homeostatic gene regulator in macrophages^3^, showed dampened upregulation by hypoxia in HFD-BMDMs (Fig. 5c). Among the homeostatic genes that can be enriched by histone lactylation^3^, *Arg1* and *Crem* gene expression levels were upregulated under hypoxia in ND-BMDMs, while the hypoxia-induced response was dampened in HFD-BMDMs (Fig. 5d, Supplementary Fig. 9). Arg1 protein levels are minimally affected (Supplementary Fig. 8). Other downstream genes on histone lactylation such as *Hsd11b1*, *Trem1*, and *Gpr18* did not follow the trend of histone lactylation in our BMDMs (Supplementary Fig. 9). As hypoxia-induced metabolic adaptation hallmark reduced oxidative stress^20^, we measured oxidative stress levels in BMDMs after hypoxia treatment using CellROX indicator. Indeed, oxidative stress levels were decreased in ND-BMDMs, whereas the decrease was impaired in HFD-BMDMs (Fig. 5d). In summary, metabolic changes under hypoxia, the induction of glycolysis-lactate accumulation, and a reduction of cellular oxidative stress levels are not properly regulated in HFD-BMDMs.

### Obesity-impaired metabolic changes under hypoxia have a minimum impact on inflammatory cytokine gene expressions

As the metabolic state is associated with macrophage polarization characterized by inflammatory cytokines ^27^, we studied inflammatory gene inductions by hypoxia or prototypical endotoxin such as lipopolysaccharide (LPS) stimulation. Hypoxia-induced *Il1b* expression in ND-BMDMs, while overall expression level of inflammatory cytokines including *Il1b*, *Il6* and *Tnfα* were not dysregulated in HFD after hypoxia treatment up to 24 hours (Supplementary Fig. 10a–d). Moreover, HFD-BMDMs had comparable cytokine expression levels with ND-BMDMs in response to LPS treatment (Supplementary Fig. 10e,f). Taken together, obesity-impaired metabolic changes under hypoxia have a minimum impact on inflammatory cytokine gene expressions.

### Obesity-impaired metabolic changes under hypoxia lead to reduced dying cell clearance in macrophages

Another macrophage function associated with the metabolic state is dying cell clearance, in which the engulfment of dying cells, the digestion of the engulfed cellular material, and energy production for running the process depend on cellular metabolic pathways including glycolysis ^18, 19, 28^, fatty acid synthesis ^29^ and glutamine utilization ^30^, as well studied in efferocytosis (phagocytotic apoptotic cell clearance). To assess overall efferocytosis, we first used apoptotic Jurkat T cells loaded with pH-sensitive dye, by which the components of apoptotic cells emit fluorescence in low-pH phagolysosomes. Live cell imaging shows increased efferocytosis activity after prolonged hypoxia (24 h) in ND-BMDM and a reduction of hypoxia-induced efferocytosis in HFD-BMDMs (Fig. 6a,b).

**Figure 6:**
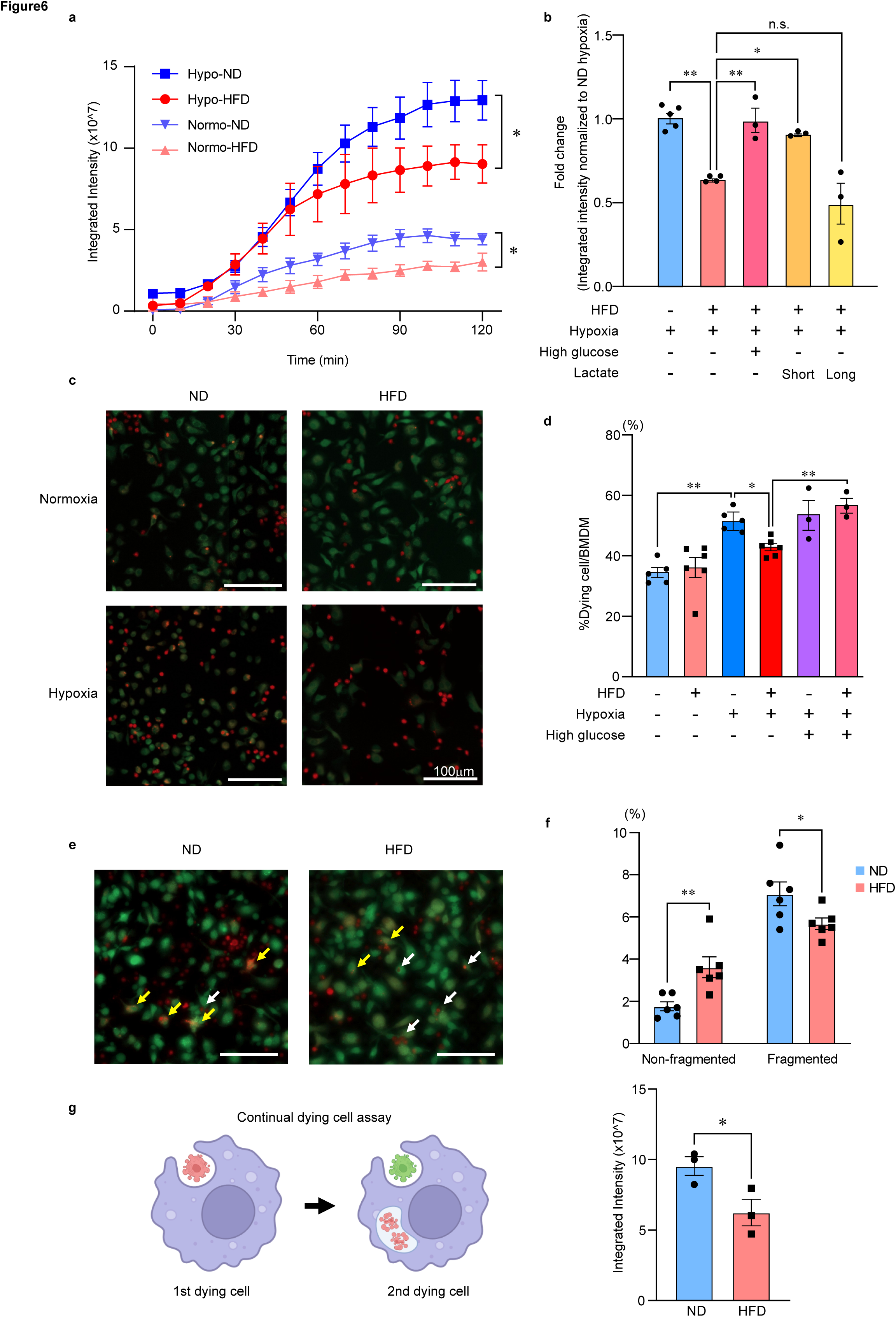
Obesity impaired dying cell clearance under hypoxic condition. a. The kinetics of dying cell clearance by ND– and HFD-BMDMs treated with or without prolonged hypoxia. n=3 biological replicates; statistical significance was determined by two-way ANOVA with Turkey’s HSD post hoc test. * p<0.05. b. Maturation of dying cell clearance evaluated by the signal intensity of pHrodo Green-labeled BMDMs was quantified relative to ND-BMDMs under hypoxia with or without high glucose (25mM). Lactate (25 mM) was treated for 24 hours before the assay (long) or only during the assay (short). n=3-5 biological replicates; statistical significance was determined by one-way ANOVA with Turkey’s HSD post hoc test. * p<0.05. c. Representative images of dying cell clearance assay with or without prolonged hypoxia treatment. CellTracker Red-labeled apoptotic neutrophils (red) were added to CSFE-labeled BMDMs (green) at a 5:1 ratio and co-cultured for 1 hour. The scale bar indicates 100µm. d. The number of CellTracker Red^+^CSFE^+^BMDMs (green and red) was quantified relative to CSFE^+^BMDMs (green only) under hypoxia with or without high glucose (25 mM). n=3-5 biological replicates; statistical significance was determined by one-way ANOVA with Turkey’s HSD post hoc test. * p<0.05. e. Representative images of non-fragmented/fragmented CellTracker Red-labeled dying cells (red) in CSFE-labeled BMDMs (green). White arrows indicate non-fragmented dying cells in BMDMs, and yellow arrows indicate fragmented dying cells in BMDMs. Scale bars indicate 100µm. f. The number of non-fragmented and fragmented CellTracker Red^+^CSFE^+^BMDMs (green and red) was quantified relative to CSFE^+^BMDMs (green only) under prolonged hypoxia. n=6 biological replicates; statistical significance was determined by Student’s t-test. g. Continual dying cell clearance was assessed after co-culture of BMDMs and dying Jurkat cells labeled with CellVue Claret Far Red (red) for 45min followed by rinsing, 2 hour-resting period, and 2^nd^ incubation of dying cell Jurkat cells labeled with pHrodo Green (green). [Left] Schematic illustrating the protocol of continual dying cell clearance. [Right] Maturation of continual dying cell clearance evaluated by the signal intensity of pHrodo Green in pHrodo Green^+^CellVue Claret^+^ BMDMs. n=3 biological replicates; statistical significance was determined by Student’s t-test. All values are means ± SEM; * p<0.05, * * p<0.01; n.s, not significant.

Next, we studied the engulfment and digestion of apoptotic neutrophils using a non-pH-sensitive dye, CellTracker Red CMTPX. Hypoxia increases the engulfment efferocytosis in ND-BMDMs, while HFD decreased efferocytosis after hypoxia treatment (Fig. 6c,d). In HFD-BMDM, macrophages with non-fragmented apoptotic neutrophils, which indicates the early phase of efferocytosis were increased, and macrophages with fragmented apoptotic neutrophils, which indicates the later phase of efferocytosis were decreased compared to ND-BMDM (Fig. 6e,f). When HFD-BMDM were cultured in high-glucose media, decreased efferocytosis capacity after hypoxia priming was rescued (Fig. 6d). Similarly, lactate supplements during efferocytosis increased efferocytosis, supporting the idea that increased lactate levels during efferocytosis promote continual efferocytosis ^19^. Indeed, we found that HFD-BMDMs have lower continual efferocytosis compared with ND-BMDMs when both BMDMs are primed under hypoxia for 24 hours (Fig. 6g,h). Finally, we tested efferocytosis in dendritic cells (DCs) grown from the BM under stimulation of granulocyte-macrophage colony-stimulating factor (GM-CSF) and did not observe differences in efferocytosis capacity between ND– and HFD-derived DCs, while hypoxia-priming increased efferocytosis (Supplementary Fig. 13). Thus, obesity-impaired metabolic changes under hypoxia lead to reduced dying cell clearance in macrophages, and the impairment quick digestion of apoptotic cells is due to reduced glucose utilization and lactate accumulation.

### Obesity induces long-term memory in hematopoietic stem cells regulating dying cell clearance capacity in differentiated BMDMs, which can be reversed by inhibiting oxidative stress in HSCs

BMDMs are grown from macrophage progenitors including most primitive hematopoietic stem cells (HSCs), which express the receptor for M-CSF. Obesity can induce long-term memory in HSCs in part through oxidative stress as observed in the similar mouse model of obesity ^31^. We confirmed that overall oxidative stress levels are increased in HSCs from HFD mice (Fig. 7a). As cyclosporin A (CSA) has been shown to reduce mitochondrial permeability transition pore-mediated oxidative stress in HSCs, we tested the effect of CSA on oxidative stress levels in HSCs from HFD mice. CSA treatment upon the harvest of BM cells can reduce mitochondria oxidative stress in HSCs from HFD mice (Fig. 7a). Measured by MitoPY1 dye with mitoTEMPO treatment, CSA likely reduced mitochondria-derived reactive oxygen species specifically in HSCs from HFD mice (Fig. 7b). Next, to determine how CSA-treatment impacts on long-term functions of HSCs, we performed serial competitive transplantation experiment. We particularly focused on dying cell clearance after hypoxia priming, which is impaired in HFD-BMDMs. We grew BMDMs after two sessions of BM transplantation into lean sub-lethally irradiated recipients (Fig. 7c). As we used EGFP mice as sources of the HSC population, which are treated with or without CSA, we were able to analyze separately the donor HSC-derived BMDMs. We found that dying cell clearance capacity is consistently reduced in HFD-HSC-derived BMDMs under hypoxia-priming (Fig. 7d). CSA treatment in HSCs during isolation from HFD mice is sufficient to rescue this obesity-induced reduction of dying cell clearance (Fig. 7 d). In the competitive bone marrow transplantation of HSCs, obesity-induced long-term memory reflects the impaired dying cell clearance in differentiated BMDMs, which is rescued by inhibiting oxidative stress in HSPCs.

**Figure 7:**
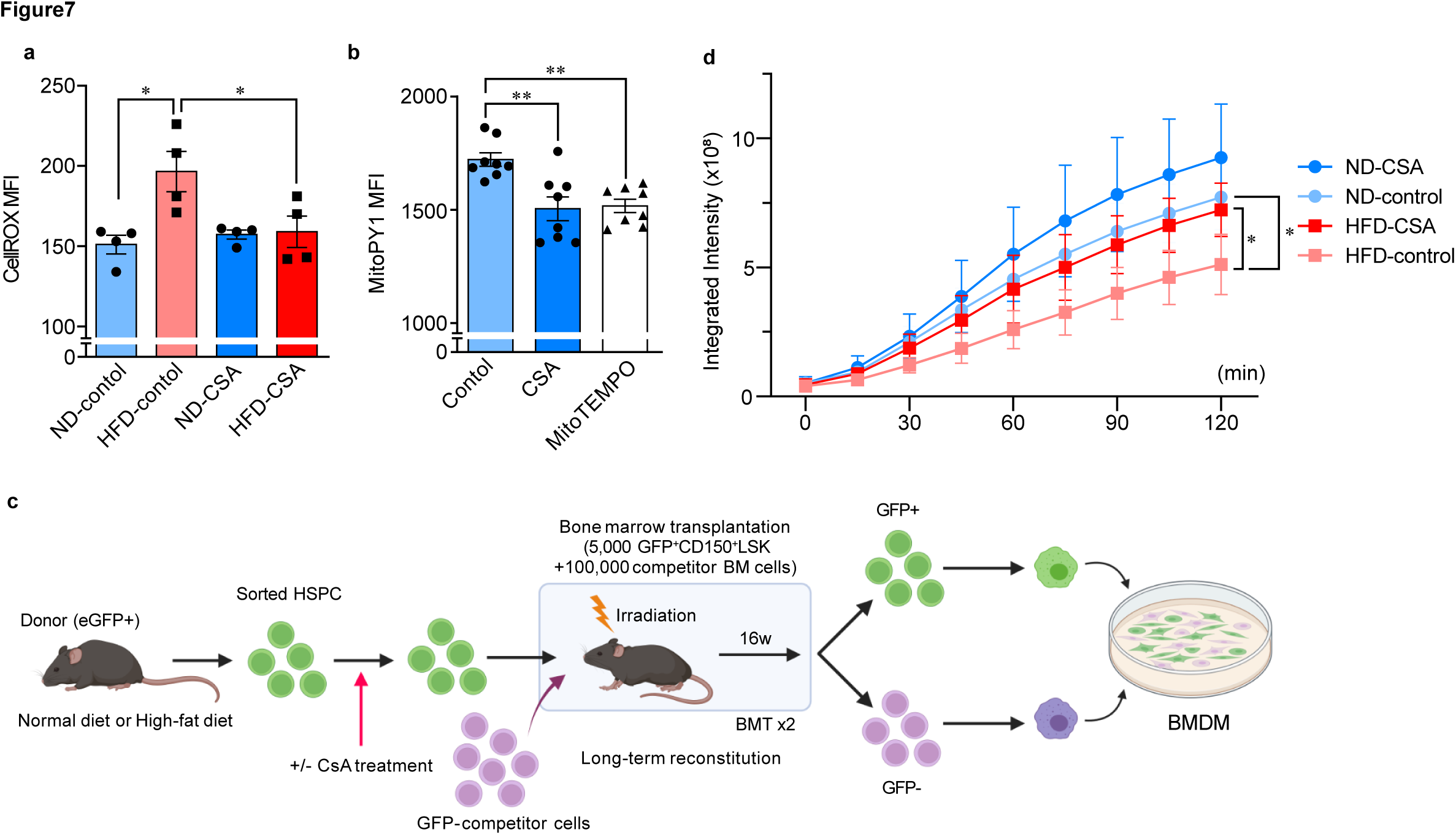
Obesity-induced long-term memory in HSPCs, which was reversed by inhibiting oxidative stress. a. Oxidative stress level in HSCs measured by CellROX Green. The measured CellROX Green signal intensity was normalized with mean fluorescence intensity (MFI). HFD increased the oxidative stress level in HSCs, which was reversed by cyclosporin A (CSA) treatment. n=4 biological replicates; statistical significance was determined by one-way ANOVA with Turkey’s HSD post hoc test. b. Oxidative stress level in mitochondria of HSCs measured by MitoPY1. The measured MitoPY1 signal intensity was normalized with MFI. CSA treatment decreased the oxidative stress level in HSCs. A mitochondria-targeted antioxidant, MitoTEMPO was used as a positive control. n=8 biological replicates; statistical significance was determined by one-way ANOVA with Turkey’s HSD post hoc test. c. Experimental scheme for serial bone marrow transplantation assay. EGFP+ HSPCs were injected into irradiated recipient mice. Peripheral blood was collected 8 and 16 weeks after transplantation for flow cytometry analysis. BMDMs were isolated 16 weeks after 2^nd^ transplantation for dying cell clearance assay. d. The kinetics of dying cell clearance by EGFP^+^BMDMs with/without CSA treatment. n=3-5 biological replicates; statistical significance was determined by two-way ANOVA with Turkey’s HSD post hoc test. All values are means ± SEM: * p<0.05, * * p<0.01.

## Discussion

In the current study, we define a mechanism by which macrophage phenotypic transition under hypoxia is regulated through chromatin response by H3K4me3 modification. Specifically, in cultured macrophages in response to hypoxia, rapidly upregulated H3K4me3 peaks are localized at promoters of the genes regulating metabolic pathways and are associated with transcriptional activation under prolonged hypoxia rather than immediate active transcription. A previous study showed rapid H3K4me3 upregulation in cultured Hela cells ^15^, however, we did not find a large overlap in the genome locations of the hypoxia-induced H3K4me3 peaks between our bone marrow-derived macrophages and Hela cells—only 7 out of 448 genes corresponding to hypoxia-induced H3K4me3 peaks in the macrophages are overlapped with the hypoxia-induced H3K4me3 peaks identified in Hela cells ^15^ (Supplementary Fig. 11). In addition, in Hela cells, H3K4me3 upregulated peaks are associated with active transcription, while the gain of H3K4me3 in macrophages likely indicates the poised chromatin for later transcriptional activation (Fig. 3). These suggest that the role of H3K4me3 response to hypoxia in transcription is context and/or cell type dependent.

We found that the broad range of impairment in the H3K4me3 response to hypoxia in obese macrophages derived from the bone marrow of HFD mice. The impairment spans from the protein levels of H3K4me3 (Fig. 1) to the genomic locations of hypoxia-modified (both upregulated and downregulated) H3K4me3 (Fig. 2). Our ChIP-seq results revealed less hypoxia-upregulated H3K4me3 peaks and more –downregulated peaks in HFD-BMDMs. Moreover, between ND-BMDMs and HFD-BMDMs, the overlaps of the hypoxia-modified H3K4me3 peaks are small — only 24 out of 330 upregulated peaks found in ND-BMDMs (Fig. 2f). The mechanism by which obesity leads to these broad dysregulations in the H3K4me3 response to hypoxia remains to be studied. KDM5 families are histone demethylases that regulate H3K4me3 levels. We did not find differences in the gene expressions of KDM5 families between BMDMs derived from ND and HFD mice (Supplementary Fig. 12). KDM5A has been identified as a direct oxygen sensor for H3K4me3, but its total protein levels are not changed (Supplemental Fig. 12), while KDM5A’s H3K4 demethylase activity can be regulated by other mechanisms such as its translocation to nucleus ^32^. Whether obesity dysregulates H3K4me3 response in a KDM5-dependent manner or through other systems such as methyltransferases will be formally determined.

Our results show some overlaps in the gene set and the individual gene level, between rapidly upregulated H3K4me3 peaks and upregulated transcription under prolonged hypoxia (Fig. 3). While we interpret this as such the gained H3K4me3 peaks by hypoxia induce the poised chromatin state for later activation, the role of H3K4me3 in transcription can be more complex ^33^. For example, a model suggests that H3K4me3, rather than initiate, promotes the elongation of RNA during transcription ^34^. Further investigation is needed to elucidate the impact of obesity-dysregulated H3K4me3 response to hypoxia on transcription.

We found that hypoxia changes metabolic status in macrophages, resulting in lactate accumulation (Fig. 5a) likely by increasing glycolysis^35^ and this metabolic change is blunted in obese macrophages. A metabolomics study of hypoxia-exposed murine macrophages showed that glucose is utilized to accumulate reduced nicotinamide adenine dinucleotide phosphate (NADPH) through a noncanonical pentose phosphate pathway (PPP) loop, which helps to reduce oxidative stress levels^20^. In line with this, we observed hypoxia-decreased oxidative stress under hypoxia in normal macrophages and impairment of this response in obese macrophages (Fig. 5d), highlighting impaired glucose utilization in obese macrophages under hypoxia. This notion is supported by the result showing glucose supplement can rescue the defective dying cell engulfment, which is a glucose-dependent process ^18^, in obese macrophages (Fig. 6b). Mechanistically, we demonstrated that a key enzyme in glycolysis, *Aldoa* is specifically downregulated in our obese macrophages after hypoxia treatment (Fig. 5a). However, the discrepancy of transcript levels and metabolic enzyme activity has been notified in macrophages under prolonged hypoxia, suggesting a complex mechanism of hypoxia-induced metabolic changes such as poised state for translation^20^. Overall, we identified rapidly-modified H3K4me3 marks on many metabolic pathway genes (both upregulated and downregulated) in normal macrophages as a potential direct oxygen-sensing response ^15^, and their dysregulation in obese macrophages. This impaired oxygen-sensing chromatin response likely influences downstream metabolic pathways beyond glucose utilization since the crosstalk between metabolites and epigenetic modification ^36^. Likewise, histone lactylation and its downstream target *Arg1* expression under hypoxia are impaired in obese macrophages (Fig. 5b and c).

Dying cell clearance including efferocytosis promotes the resolution of inflammation^37^ and is a critical step for wound healing and tissue repair^38^. Efferocytosis is associated with macrophage polarization, in which IL-4-induced M2-like macrophages have a higher capacity of this process^39^, while efferocytosis induces glycolysis by promoting *Slc2a* and *Pfkfb2* expression that has been associated with pro-inflammatory M1-like activation ^18, 19^. In macrophages primed by prolonged hypoxia (for 7 days), the upregulation of glycolytic genes and reduced oxidative stress favor efferocytosis capacity ^20^. In line with these findings, our data demonstrated hypoxia-induced glycolysis and lactate accumulation, and reduced oxidative stress in normal macrophages and show the impairment of these processes in obese macrophages leads to reduced capacity of dying cell clearance (Fig. 6). Indeed, continual efferocytosis is lactate dependent ^19^.

In our hypoxia-primed macrophages (for 24 hours), we found upregulation in *Arg1*, *Aldoa* and *Crem* (Fig. 4e, 5c and Supplementary Fig. 9), but not in other efferocytosis-related genes such as *Trem1* and *Fgr* (Supplementary Fig. 9). Importantly, *Arg1* is essential for actin reorganization during engulfment of dying cells ^39^. Further investigation will be needed to determine the link between hypoxia-induced H3K4me3 peaks and transcriptional activation during efferocytosis.

Our results indicate that metabolic rewiring is dysregulated in obese macrophages. Glycolysis is associated with inflammatory macrophages^40^. Emerging evidence supports the notion that the glycolytic state is a result rather than a mediator of inflammatory polarization of macrophages since nitric oxide produced in M1-like macrophages activates glycolysis and represses oxidative phosphorylation^41^. In normal macrophages, glycolysis and subsequent lactate accumulation activate transitional mechanisms to anti-inflammatory macrophages as lactate acts in lysosomes to activate mTORC1 and HIF-2α^42^; lactate acts in mitochondria by binding to mitochondrial antiviral-signaling protein (MAVS) and inhibiting its interaction with RIG-I^43^; lactate-induced extracellular acidosis activates the transcriptional repressor cAMP-responsive element modulator (CREM), which suppresses inflammatory genes^44^. Despite the reduction of histone lactylation and its target *Crem* and *Arg1* gene expression in hypoxia-primed obese macrophages (Fig. 5c and Supplementary Fig. 9), our obese macrophages did not change antiinflammatory programming under prolonged hypoxia (Supplemental Fig. 10). Alongside recent studies^19, 28^, our study highlights the significance of lactate management in efferocytosis and its downstream events in macrophages. An *in vivo* significance of obesity-dysregulated metabolic rewiring requires further investigation with a note that enhanced glycolysis and HIF-1α activation has been documented in adipose tissue macrophages in obesity^45^, which may be a result of impaired metabolic rewiring in response to a local environment including hypoxia.

We show biochemical, metabolic, and phenotypic differences in BMDMs between obese and non-obese mice. The crosstalk between systemic metabolism and immune cell state has been implicated ^46^, and long-term immune memory can be formed in bone marrow stem cells ^47^. Indeed, in rodent obesity models with a transplantation setting, long-term functional changes have been observed at the level of hematopoietic stem cells ^31, 48, 49^. We found M-CSF, but not GM-CSF, differentiated macrophages exhibited obesity-perturbed phenotype in terms of dying cell clearance capacity (Fig. 6). Given that bone marrow-derived monocytes/macrophages contribute to tissue macrophages in many organs, there is likely a mechanistic link between dysregulated hypoxia response through chromatin and perturbed tissue macrophages in obesity. In addition, the link between hypoxia response through chromatin, metabolic rewiring, and dying cell clearance capacity, which activates the resolution mechanism, will open a new avenue to targeting chromatin response as a therapeutic strategy for obesity-associated non-resolution.

## Figure Legends

**Supplementary Figure 1:**
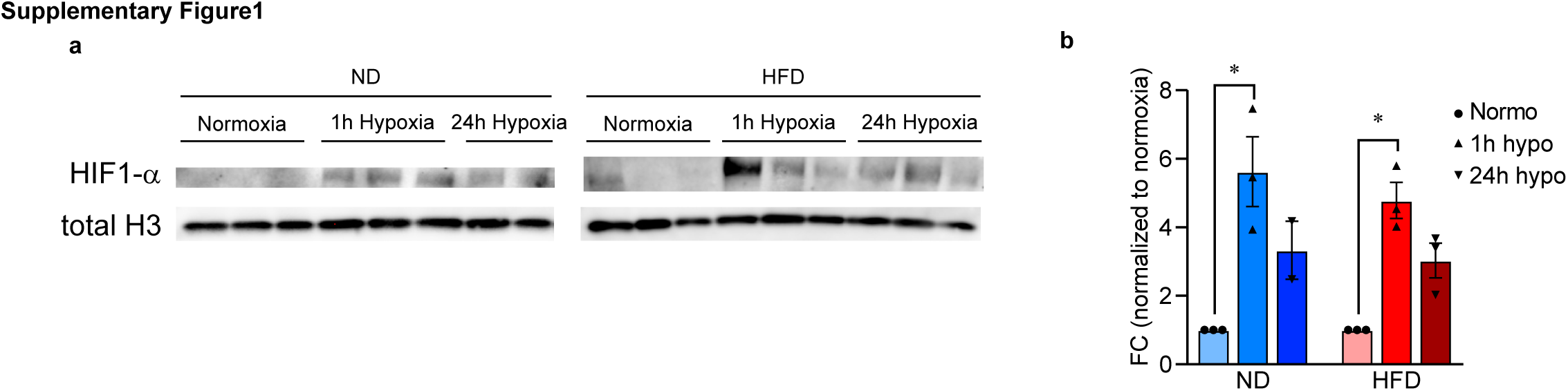
HIF1 induction after hypoxia treatment is intact in HFD-BMDMs. a. Immunoblot of cytosolic HIF1α level in BMDMs after hypoxia treatment (for 1 and 24 hours). b. The relative expressions of cytosolic HIF1α versus β-actin were quantified by band densitometry. n=3 biological replicates; statistical significance was determined by two-way ANOVA with Turkey’s HSD post hoc test. All values are means ± SEM: * p<0.05.

**Supplementary Figure 2:**
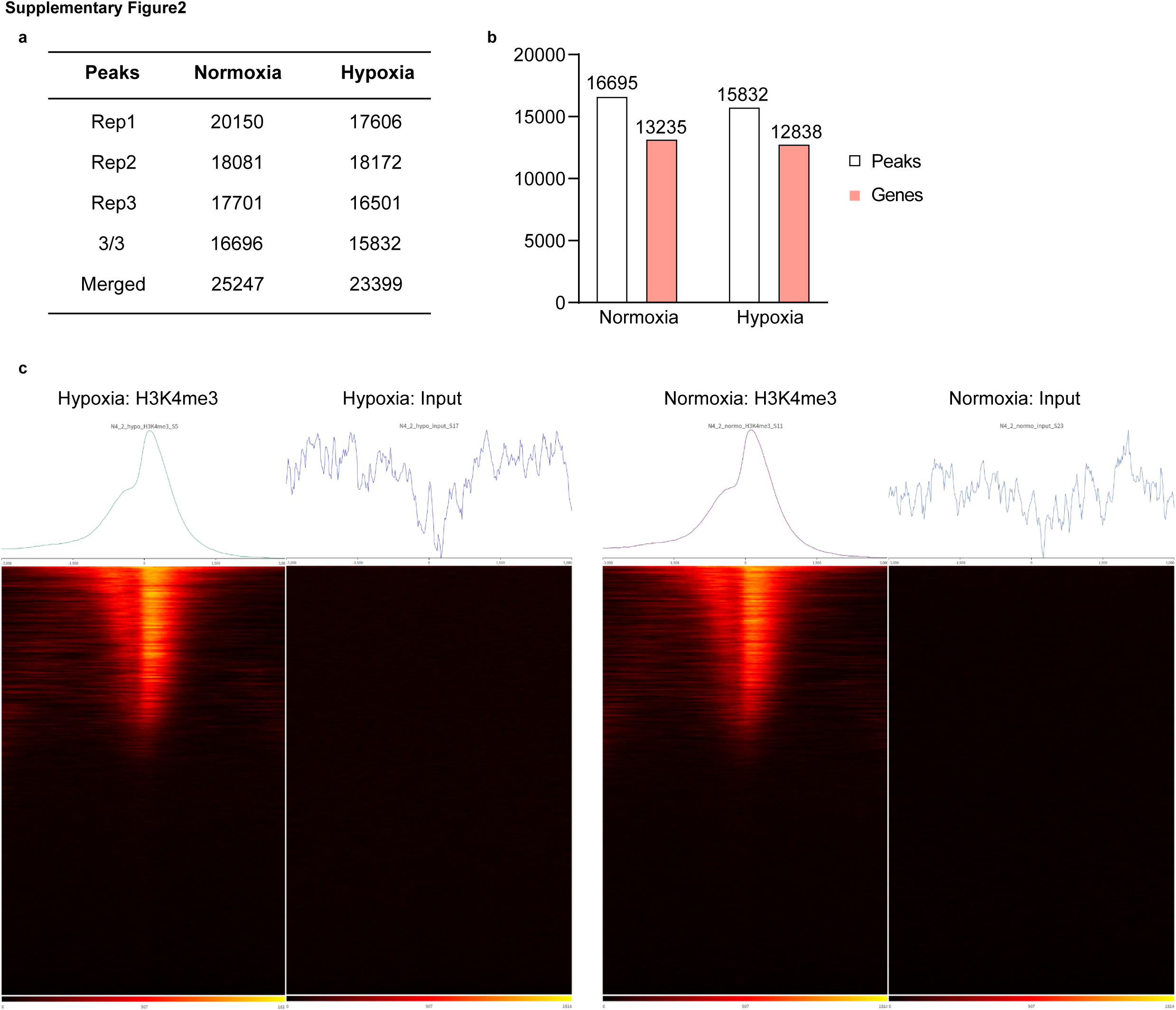
H3K4me3 ChIP-seq analysis. a. Number of peaks called in each condition, 3/3 represents peaks called all 3 replicates (high stringency peaks), and merged represents the merging data from all 3 replicates. b. High stringency peaks and genes (genes with peaks) called in each condition. c. Heatmap of CPM normalized read counts at Transcription Start Sites (TSS) of protein-coding genes for each condition.

**Supplementary Figure 3:**
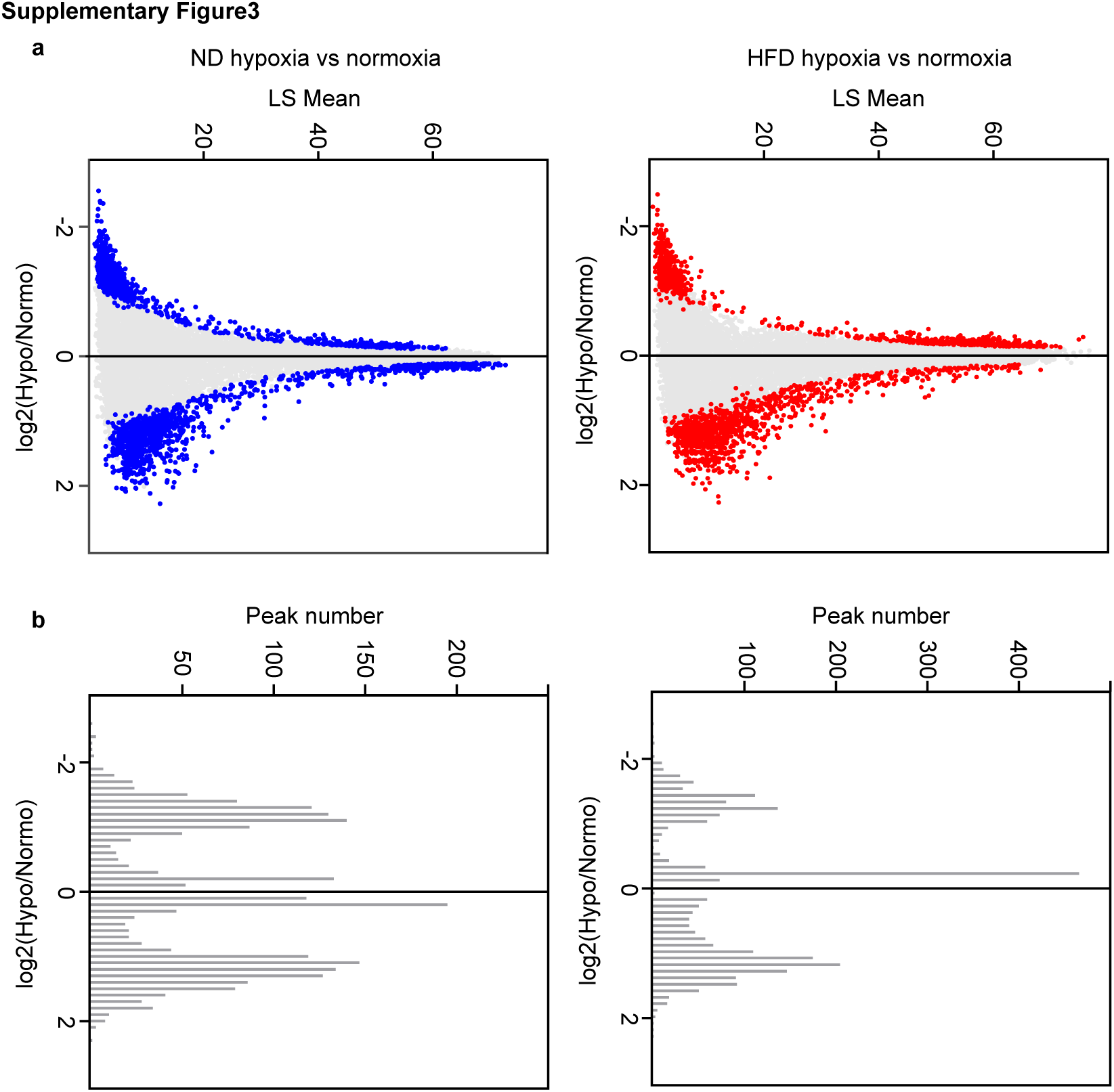
Relationship of fold change and peak height in H3K4me3 ChIP-seq. a. Relationship of fold change (y-axis) and LSmean (x-axis) in H3K4me3 peaks in response to hypoxia in ND-BMDMs (left) and HFD-BMDMs (right). b. Relationship of fold change (y-axis) and number of peaks (x-axis) in H3K4me3 peaks in response to hypoxia in ND-BMDMs (left) and HFD-BMDMs (right).

**Supplementary Figure 4:**
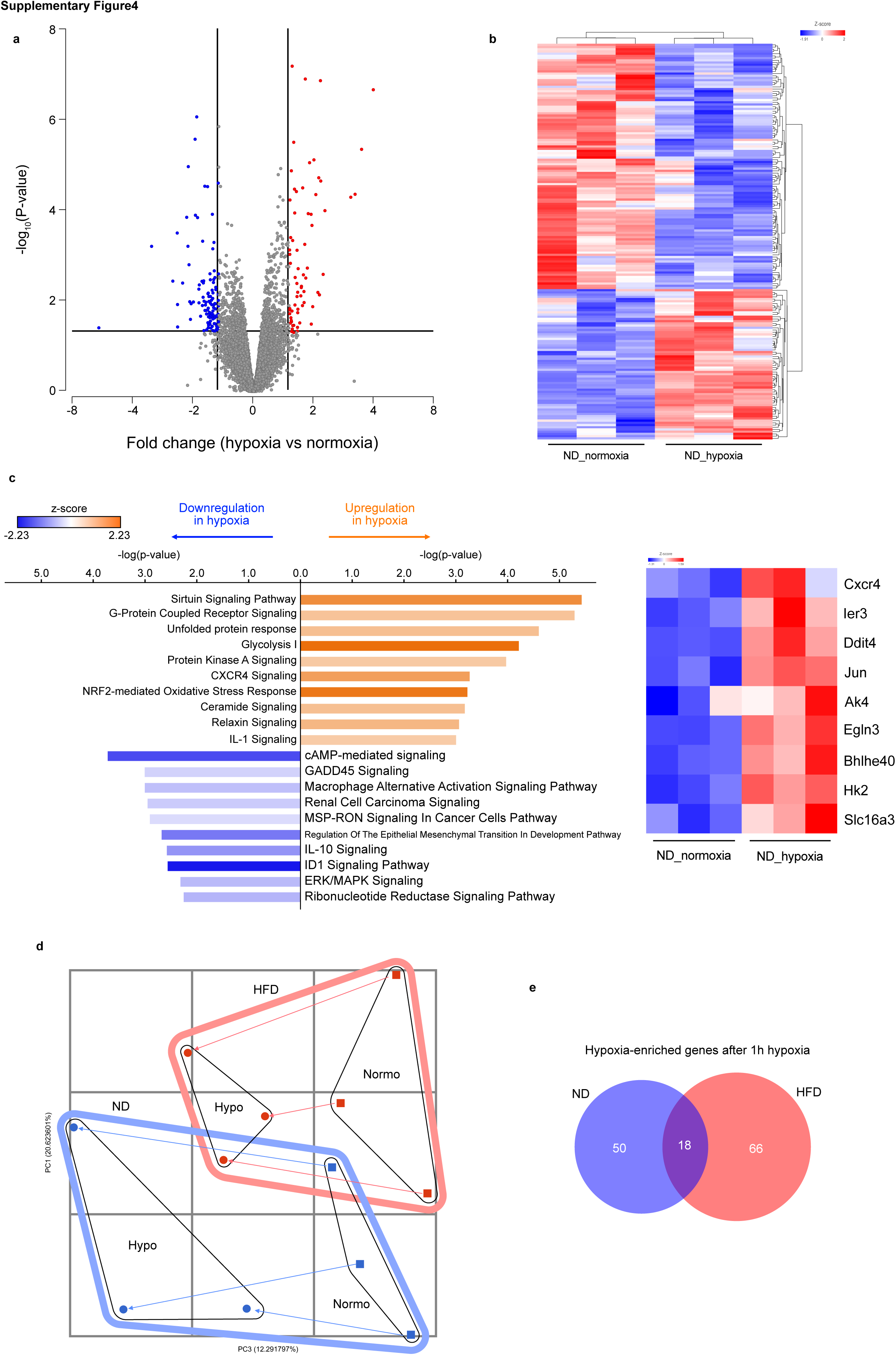
Hypoxia-induced genes in ND-BMDM identified by RNA-seq. a. Volcano plot showing differentially expressed genes after brief hypoxia (1 hour) in ND-BMDMs. After hypoxia treatment, 69 and 112 genes were upregulated and downregulated, respectively. The horizontal line indicates a p-value of 0.05, and the vertical line a fold change of ±1.5. b. Heat map showing differentially expressed genes after hypoxia treatment. There is a clear separation in expression pattern between the normoxia and hypoxia groups. c. [Left] Gene ontology (GO) analysis with ingenuity pathway analysis (IPA) revealed that hypoxia-upregulated genes were enriched in glycolysis-related pathways. [Right] Heat map showing the individual expression pattern of glycolysis-related genes. d. Principal component analysis (PCA) plot showed hypoxia altered the gene expression profile both in BM– and HFD-BMDMs. e. Overlaps of hypoxia-upregulated genes between ND– and HFD-BMDMs. n=3 biological replicates. Statistical significance was determined by GSA.

**Supplementary Figure 5:**
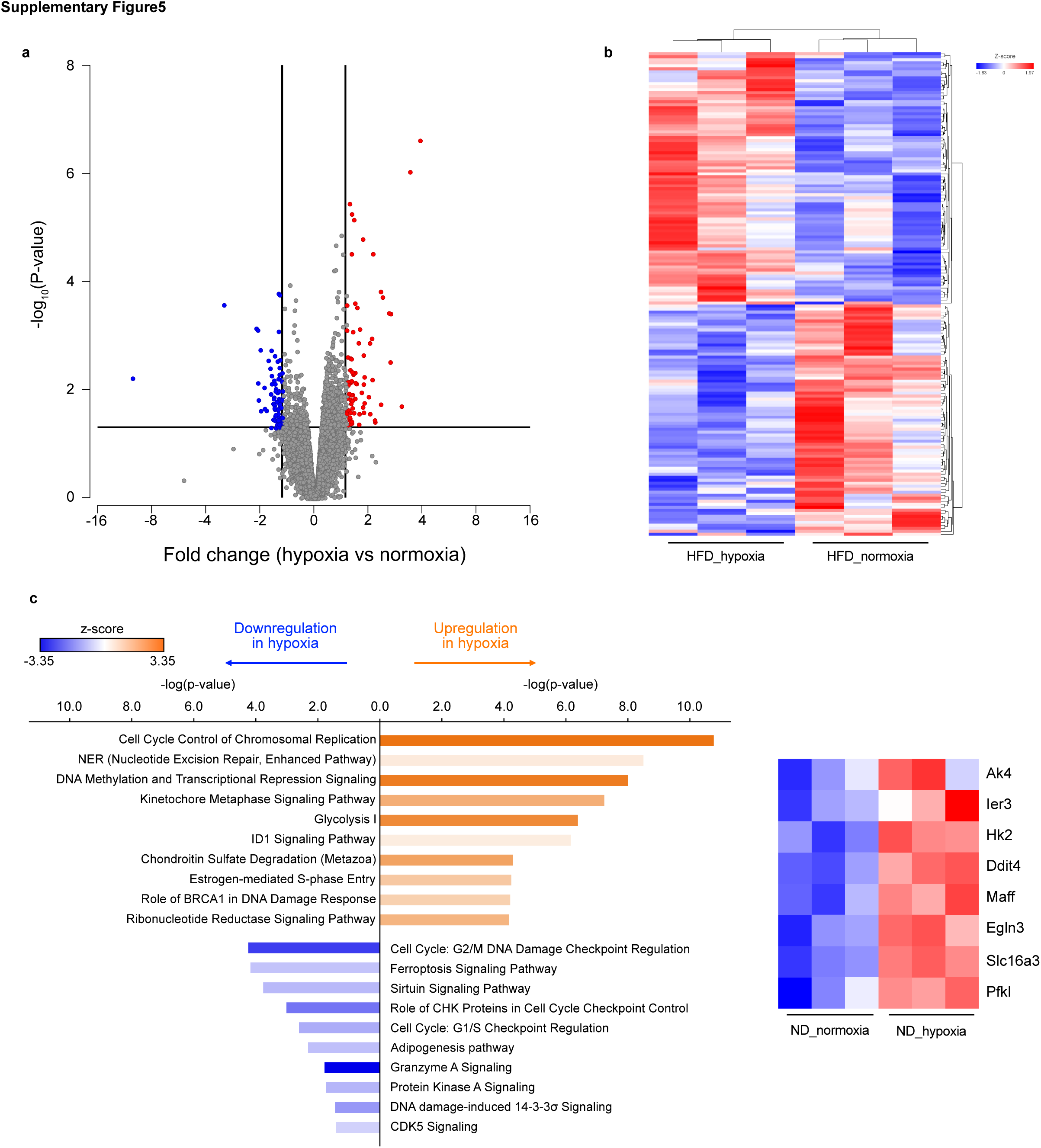
Hypoxia-induced genes in HFD-BMDM identified by RNA-seq. a. Volcano plot showing differentially expressed genes after brief hypoxia (1 hour) in HFD-BMDMs. After hypoxia treatment, 84 and 77 genes were upregulated and downregulated, respectively. The horizontal line indicates a p-value of 0.05, and the vertical line a fold change of ±1.5. b. Heat map showing differentially expressed genes after hypoxia treatment. There is a clear separation in expression pattern between the normoxia and hypoxia groups. c. [Left] Gene ontology (GO) analysis with ingenuity pathway analysis (IPA) revealed that hypoxia-upregulated genes were enriched in glycolysis-related pathways. [Right] Heat map showing the individual expression pattern of glycolysis-related genes. n=3 biological replicates. Statistical significance was determined by GSA.

**Supplementary Figure 6:**
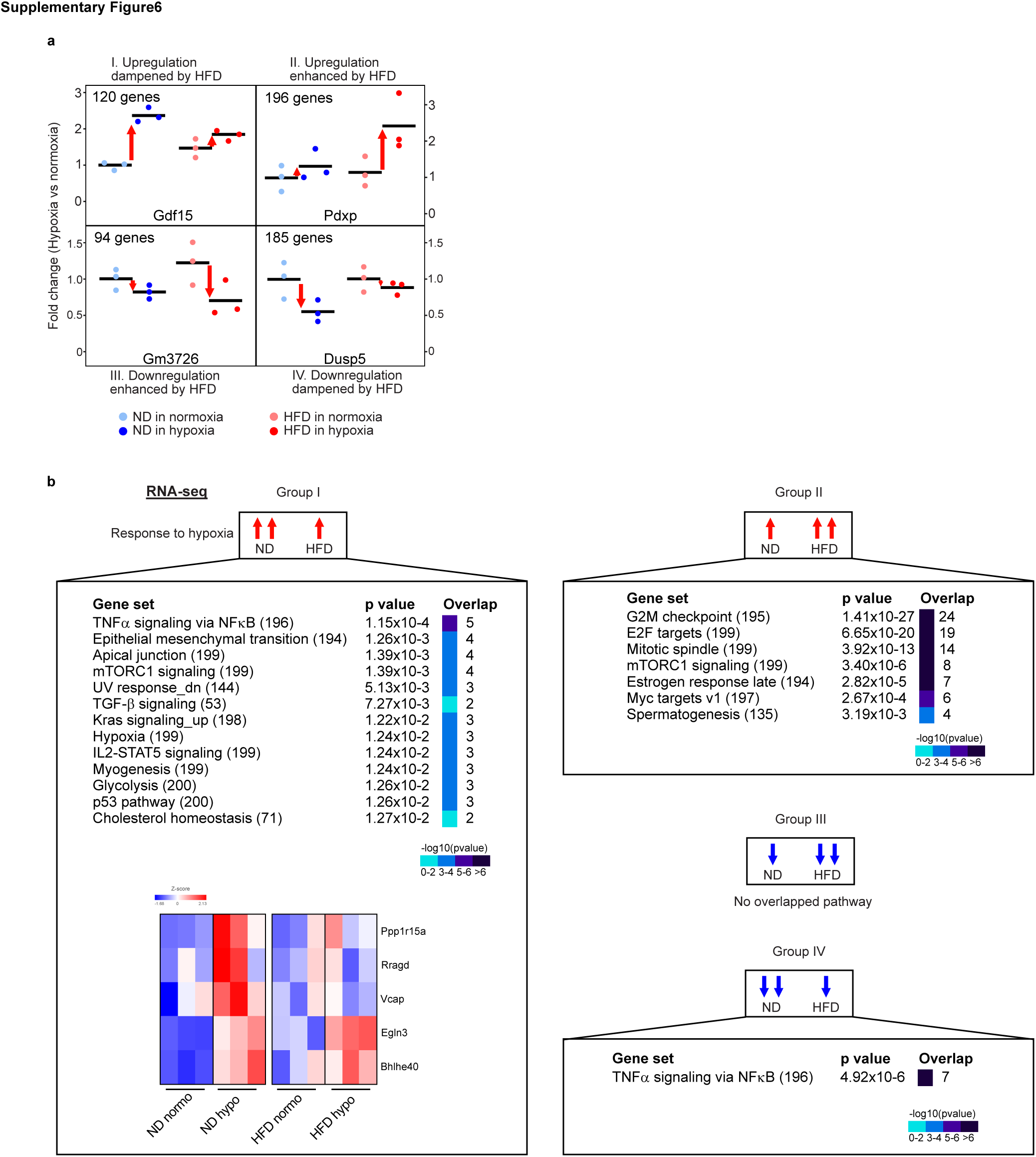
Obesity-impaired hypoxia-induced gene expression pattern in metabolic genes. a. Categorization of gene expression pattern identified by RNA-seq. I: the gene set with hypoxia-induced upregulation dampened by HFD. II: the gene set with hypoxia-induced upregulation enhanced by HFD. III: the gene set with hypoxia-induced downregulation enhanced by HFD. IV: the gene set with hypoxia-induced downregulation dampened by HFD. b. Gene ontology (GO) analysis revealed that metabolic pathways are enriched with the Group1 gene set, while other gene sets do not have an association with metabolic pathways.

**Supplementary Figure 7:**
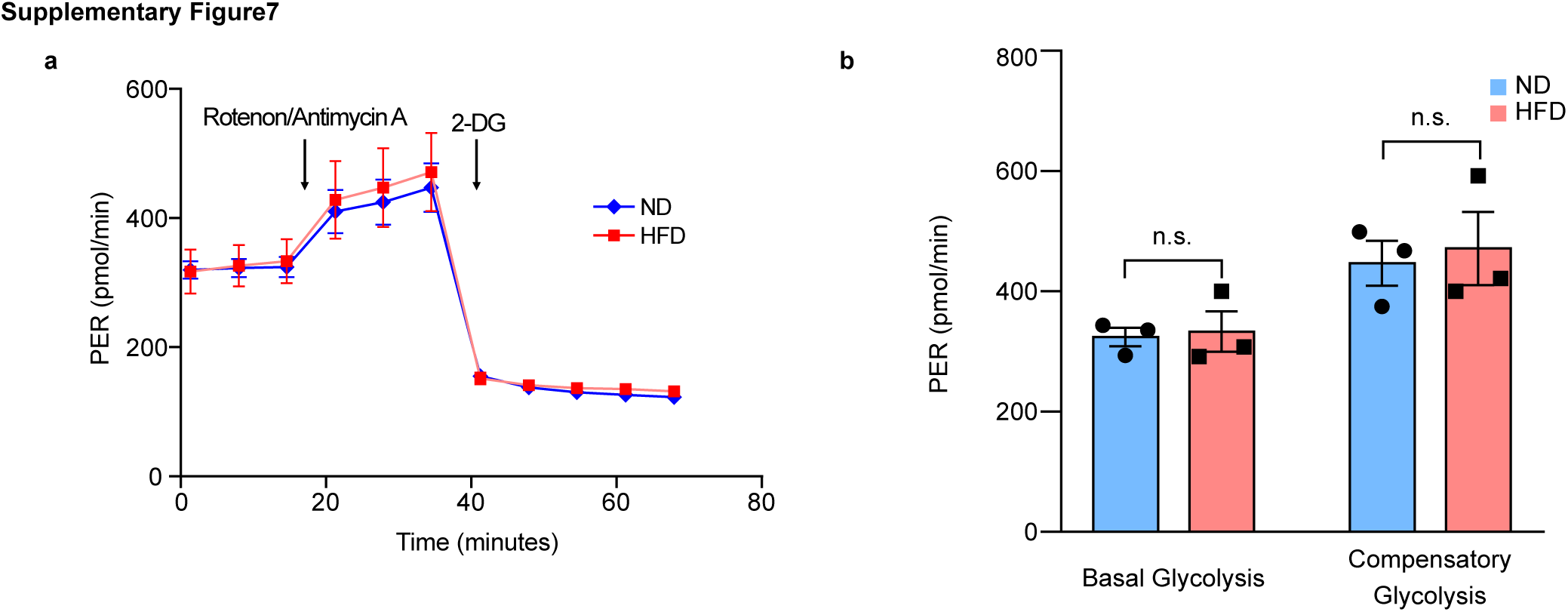
Obesity didn’t impair glycolytic capacity under normoxia. a. The glycolytic proton efflux rate (PER) of BMDMs was analyzed by Seahorse XFe96 analyzer. Rotenone and antimycin A were added to determine compensatory PER following basal measurement of PER., This was followed by the addition of 2-deoxy-D-glucose to ensure that the PER observed was caused by glycolysis. b. Basal and compensatory glycolytic PER were comparable between ND– and HFD-BMDMs. n=3 biological replicates; statistical significance was determined by Student’s t-test. All values are means ± SEM; n.s., not significant.

**Supplementary Figure 8:**
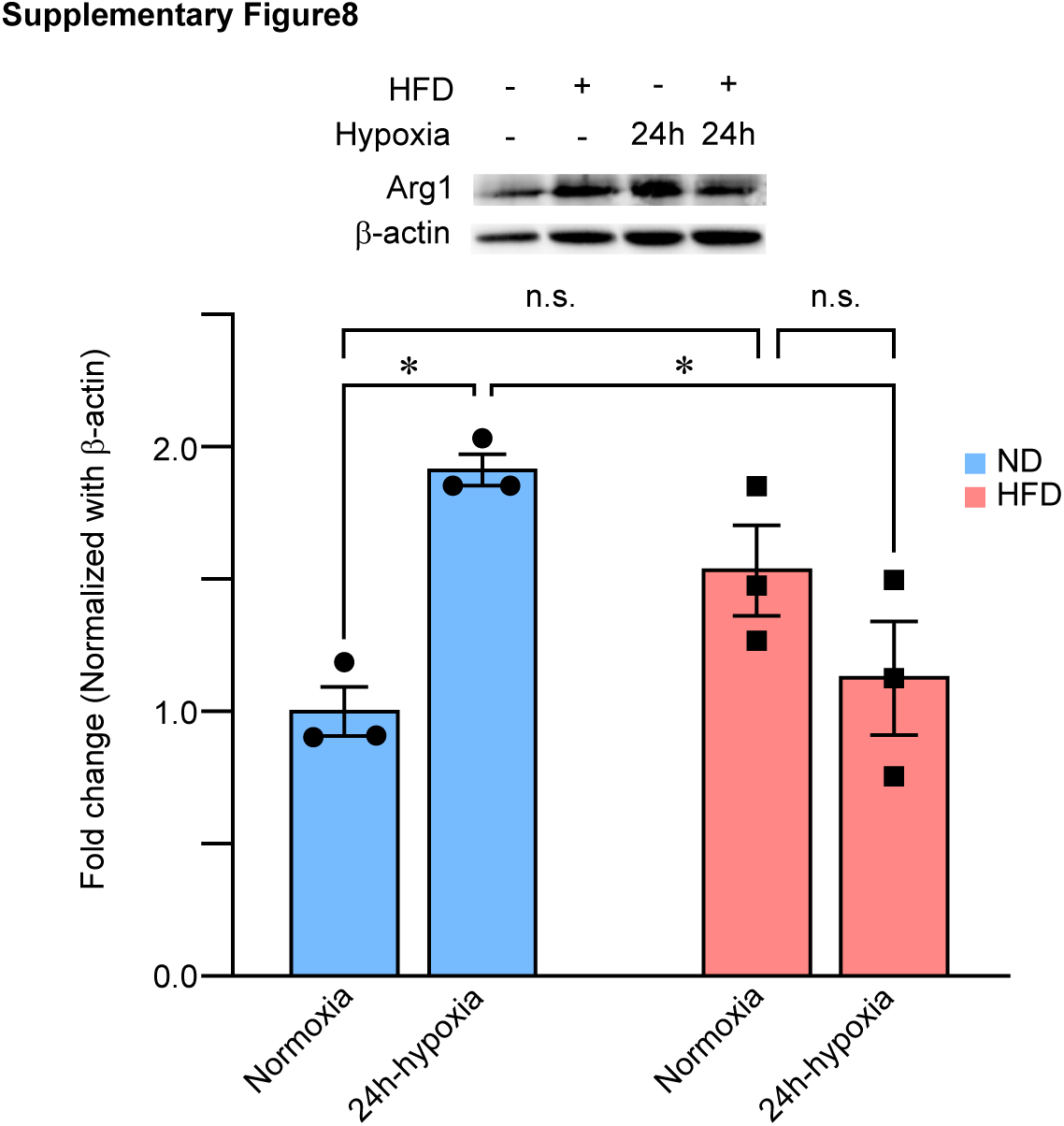
Arg1 protein expression level. [Top] Immunoblots of Arg1 from BMDMs in response to prolonged hypoxia. [Bottom] The relative amounts of Arg1 versus β-actin were quantified by densitometry. n=3 biological replicates; statistical significance was determined by two-way ANOVA with Turkey’s HSD post hoc test. All values are means ± SEM: * p<0.05; n.s., not significant.

**Supplementary Figure 9:**
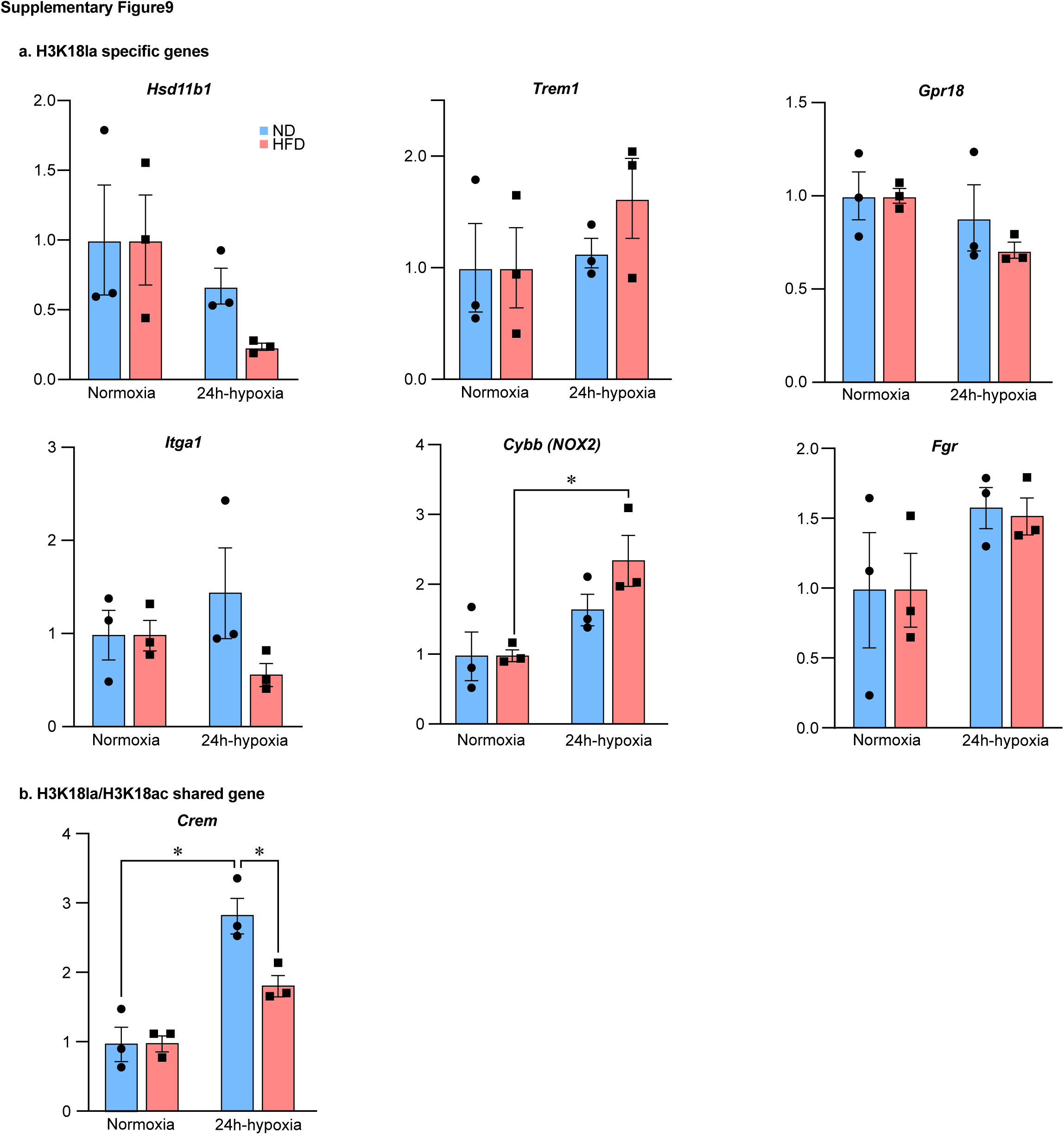
Gene expression profile of histone-lactylation regulating genes in response to prolonged hypoxia. a. Hypoxia-inducible gene expression of *Arg1* in ND-BMDMs and HFD-BMDMs. The rate of hypoxia induction was assessed by normalizing the relative expression level in hypoxia with the one in normoxia. n=3 biological replicates; statistical significance was determined by two-way ANOVA with Turkey’s HSD post hoc test. All values are means ± SEM: * p<0.05.

**Supplementary Figure 10:**
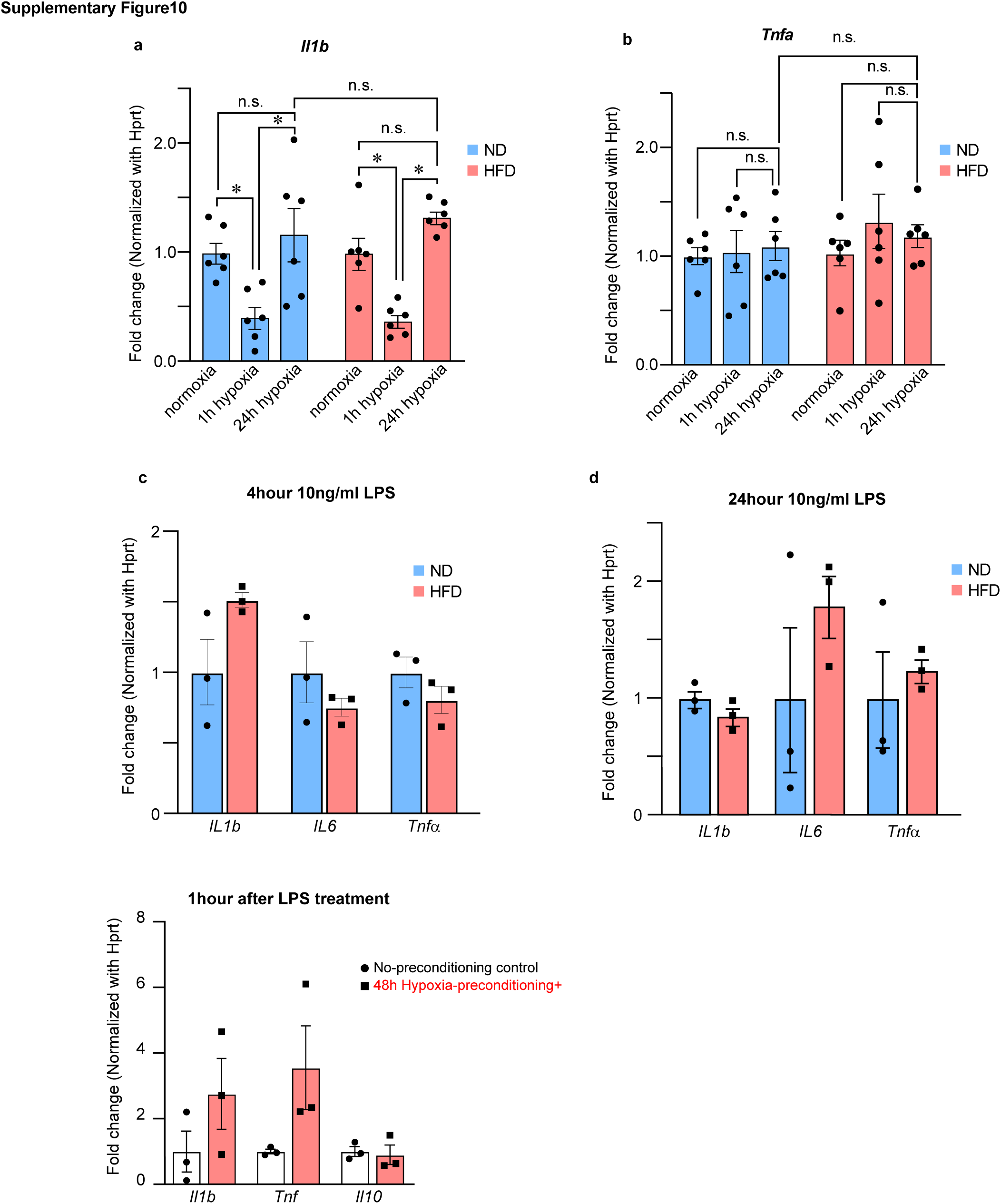
Gene expression profile of cytokines in response to hypoxia and LPS. a-b. Cytokine expression levels in ND– and HFD-BMDMs were analyzed by qPCR after a brief and prolonged hypoxia. Relative expression of cytokine genes versus *Hprt* HFD-BMDMs to ND-BMDMs. n=6 biological replicates; statistical significance was determined by two-way ANOVA with Turkey’s HSD post hoc test. c-d. Cytokine expression levels in ND– and HFD-BMDMs were analyzed by qPCR after LPS stimulation for 4 hours (c) and 24 hours (d). Relative expression of cytokine genes versus *Hprt* HFD-BMDMs to ND-BMDMs. n=3 biological replicates; statistical significance was determined by two-way ANOVA with Turkey’s HSD post hoc test. All values are means ± SEM: * p<0.05; n.s., not significant.

**Supplementary Figure 11:**
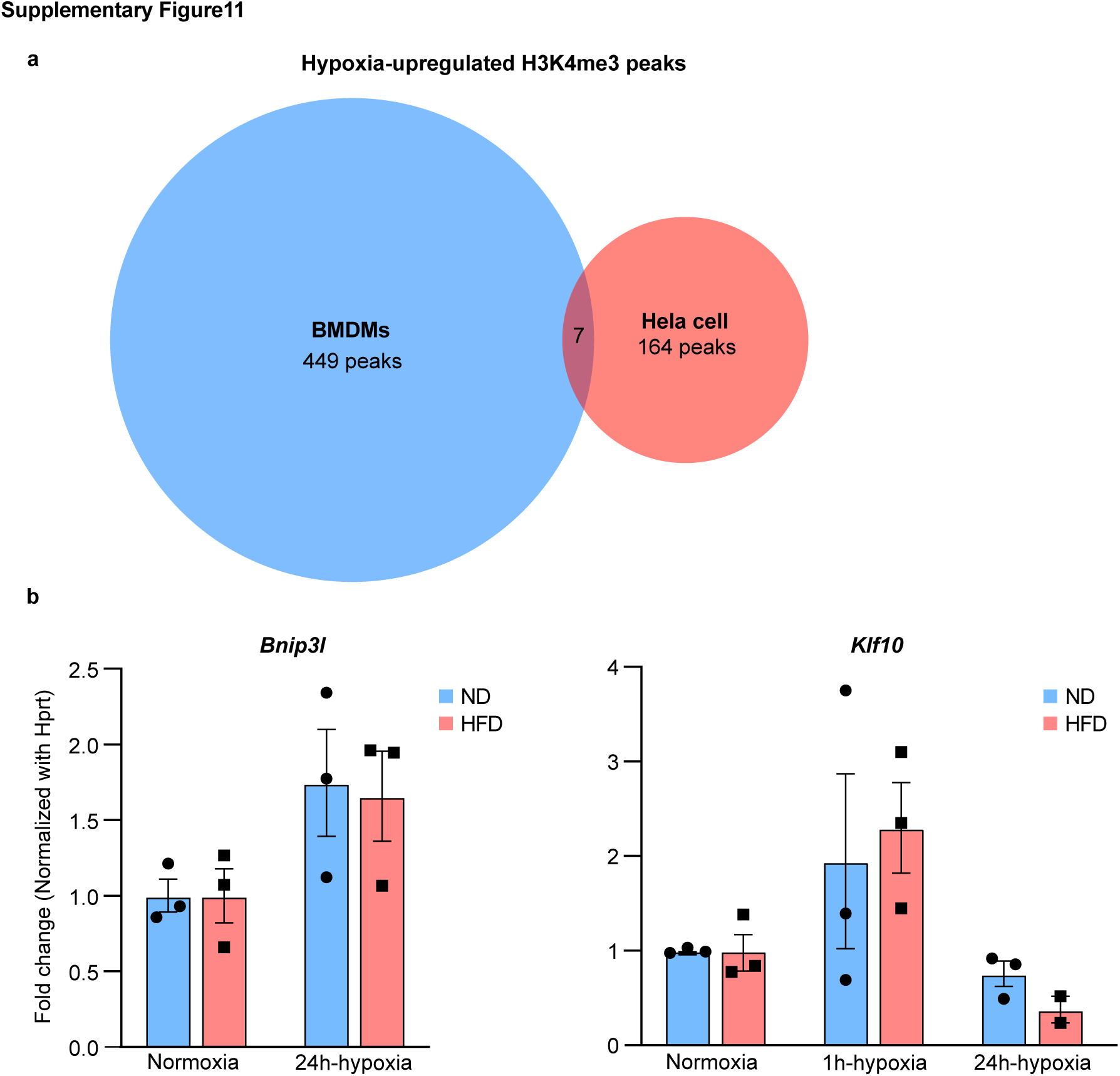
Overlaps of hypoxia-upregulated genes between BMDMs and Hela cells. a. Overlaps of hypoxia-upregulated genes between BMDMs and Hela cells illustrated with a Venn diagram. b. The gene expression levels of the Kdm5a downstream target were measured by qPCR. n=3 biological replicates; statistical significance was determined by two-way ANOVA with Turkey’s HSD post hoc test. All values are means ± SEM; * p<0.05. n.s., not significant.

**Supplementary Figure 12:**
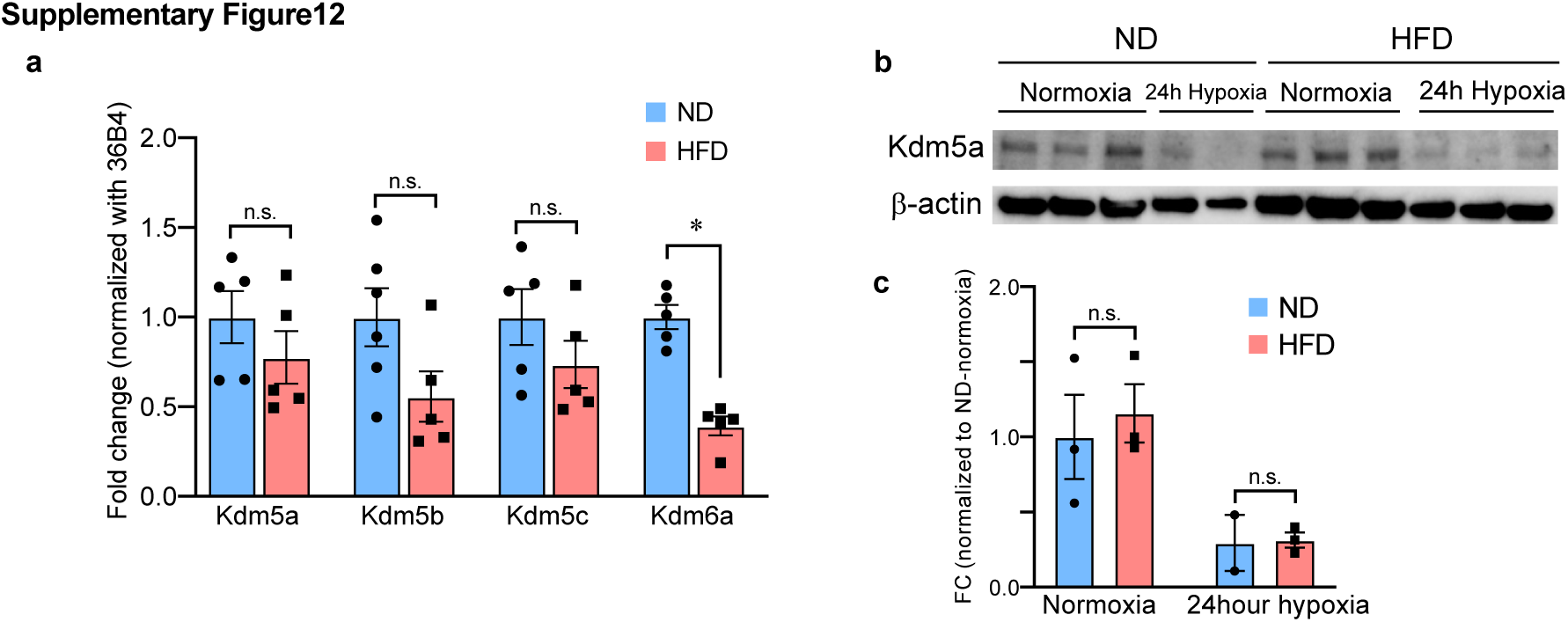
Kdm5 family expression after hypoxia treatment. a. The gene expression levels of the Kdm5 family were measured by qPCR. n=5-6 biological replicates; statistical significance was determined by Student’s t-test. b. Immunoblot of cytosolic Kdm5a level in BMDMs after 24h hypoxia. c. The relative expressions of cytosolic Kdm5a versus β-actin were quantified by band densitometry. n=3 biological replicates; statistical significance was determined by two-way ANOVA with Turkey’s HSD post hoc test. All values are means ± SEM; * p<0.05. n.s., not significant.

**Supplementary Figure 13:**
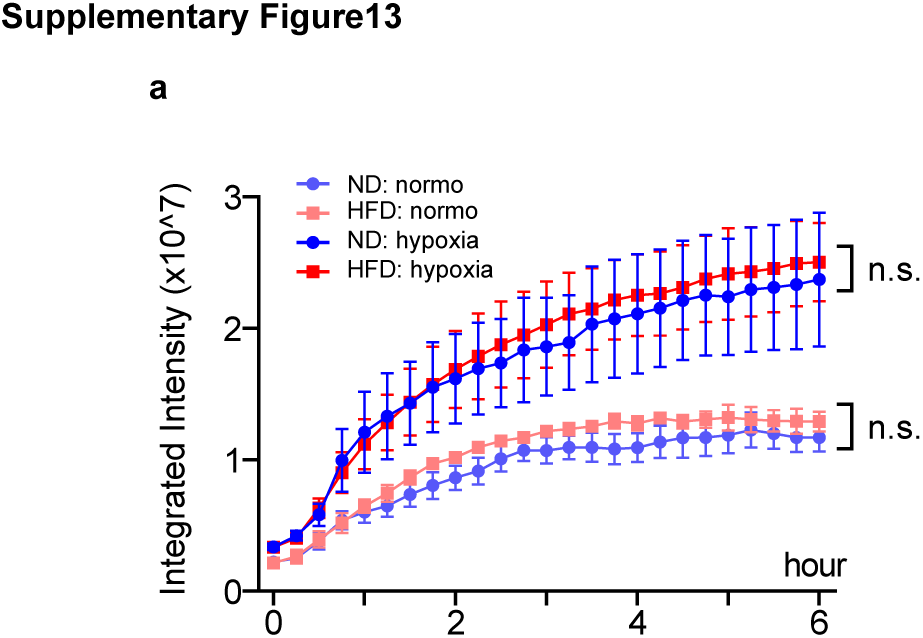
Dying cell clearance capacity in dendritic cells. a. The kinetics of dying cell clearance by ND– and HFD-dendritic cells treated with or without prolonged hypoxia. n=3 biological replicates; statistical significance was determined by two-way ANOVA with Turkey’s HSD post hoc test. All values are means ± SEM; n.s., not significant.

